# Predicting single-molecule chromatin distribution from DNA sequence

**DOI:** 10.64898/2026.10.29.762992

**Authors:** Ruoyu Wang, Junru Jin, Sebastian Pott, Jian Zhou

**Affiliations:** Section of Genetic Medicine, Department of Medicine, University of Chicago, Chicago, IL 60637, USA

## Abstract

Genome regulation unfolds through dynamic and stochastic single-molecule events. Deep learning models that read gene regulatory activities from DNA sequences have markedly advanced our ability to predict cell-population-level averages. However, they cannot capture the full spectrum of heterogeneity and dynamics at the single-molecule level, which directly reflect the underlying mechanisms of genome regulation. Here, we introduce a sequence-to-distribution (S2d) framework that directly models the entire probabilistic distribution of single-molecule chromatin states from DNA sequence alone. We used this framework to train Mola, a model of single-molecule chromatin accessibility. We harness Mola to investigate sequence-based regulatory code and mechanisms for multiple aspects of chromatin regulation at the single-molecule level. By quantifying sequence-driven nucleosome regulation, we reveal distinct modes of transcription factor engagement with nucleosomes, including a novel nucleosome-trapping factor, ZNF146. We characterized the sequence determinants governing the +1 nucleosome after transcription start sites and identified YY1 as the strongest +1 nucleosome-positioning factor. Leveraging long-context single-molecule modeling, we further discovered a shared sequence basis for the co-accessibility of regulatory elements and the co-footprinting of transcription factors. Joint prediction of DNA methylation and accessibility reveals a sequence-driven bimodal state mediated by methylation-sensitive transcription factors. Beyond static single-molecule snapshots, the S2d framework can generate pseudo-dynamic trajectories of chromatin states and propose a roadmap toward genome-wide chromatin dynamics. Moreover, we showed that Mola can predict and interpret the effects of genetic variants at the single-molecule level, revealing how sequence changes reshape distal regulatory coordination. Together, this sequence-to-distribution framework lays the foundation for understanding how genomic sequence encodes stochastic and dynamic genome regulation, and how genetic variation changes it, one molecule at a time.

## Introduction

Genome regulation is dynamic and stochastic, leading to heterogeneity of chromatin state across molecules in a population of cells^1–3^. At any given locus, nucleosomes sit at different positions on different molecules, transcription factors are bound on only a fraction of them, and a promoter and its distal enhancer are open together on some molecules but not others^1,4,5^. Together, these events shape a landscape of highly dynamic single-molecule chromatin states. Bulk chromatin measurements reduce the dynamic gene regulation process to a single profile, missing the underlying distribution of molecular states that determines how transcriptional regulation occurs.

Our capacity to measure and predict regulatory chromatin sets the pace of how much we understand the non-coding regulatory genome. Sequence-to-function models have achieved breakthroughs in predicting bulk-average levels of chromatin accessibility, TF occupancy, and transcriptional outputs from DNA sequences, and are now widely used to interpret noncoding variants^6–8^. However, all current sequence-to-function models are trained on population averages and therefore cannot understand the full spectrum of behaviors expressed by individual molecules, including heterogeneous chromatin opening, nucleosome phasing, and molecule-wide coordination between distal elements. Single-molecule chromatin mapping and modeling are imperative to ultimately understand the mechanisms of genome regulation.

Single-molecule sequencing technologies can read out long DNA molecules and directly detect DNA modifications^9^. Building on these platforms, methods such as Fiber-seq, which combines methyltransferase footprinting with long-read sequencing, enable nucleotide-resolution analysis of individual chromatin fibers, revealing single-molecule heterogeneity of chromatin accessibility, nucleosome positioning, and transcription factor occupancy^10–14^. These single-molecule data necessitate a different modeling target for the field: not a single expected average profile, but the full probability distribution of chromatin states across molecules, including their interdependencies (**Figure 1A**). A sequence model that can learn this distribution would be valuable not only as a predictor but also as a mechanistic discovery engine for the sequence-based code, bringing the vision of “*virtual chromatin*” within reach. It would enable controlled *in silico* sequence perturbations and quantitative readouts of how molecular-state distributions change in response to sequence changes.

**Figure 1.**
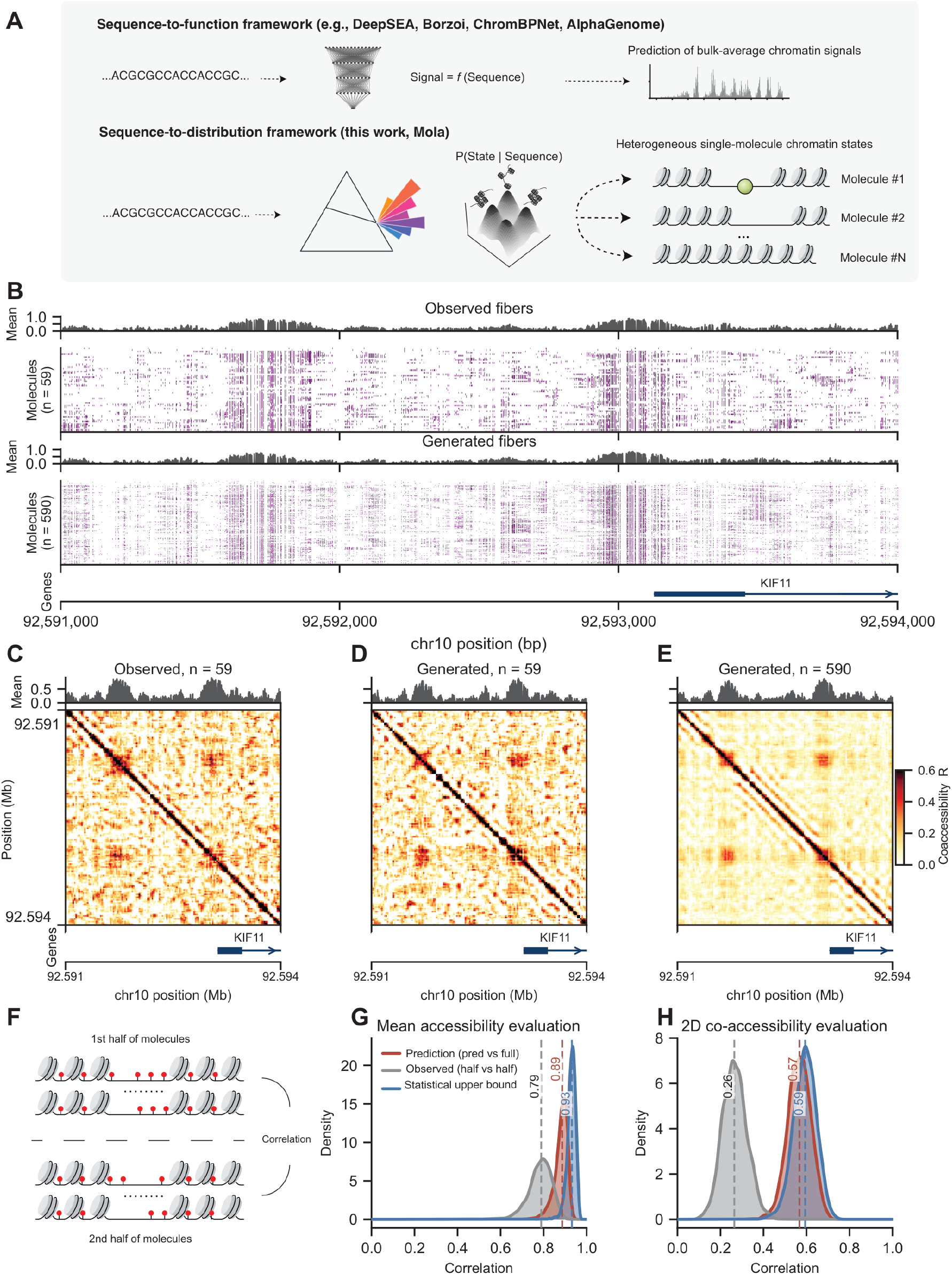
A sequence-to-distribution framework generates single-molecule chromatin states from DNA sequence. (A) Sequence-to-function (S2f) models take a DNA sequence and predict one bulk-average signal track. The sequence-to-distribution (S2d) framework instead learns the conditional distribution of chromatin states given the sequence, P(state | sequence), and draws many single-molecule states from it. Each draw is one possible chromatin state for the same sequence. (B) Observed Fiber-seq molecules (n = 59) and Mola-generated molecules (n = 590, ten times the observed number) at the KIF11 promoter on held-out chromosome 10 (3-kb view, chr10:92,591,000–92,594,000). Purple marks accessible A/T positions on each molecule. The track above each panel shows the mean accessibility across molecules. (C–E) Co-accessibility maps for the same region, defined as the Pearson correlation of per-bin accessibility across molecules, for observed molecules (C; n = 59), the same number of generated molecules (D; n = 59), and ten times more generated molecules (E; n = 590). The promoter and the upstream distal element appear as off-diagonal blocks in both observed and generated data. (F) Half-split benchmark. We split the observed molecules at each DHS into two halves and correlated the mean accessibility and co-accessibility of one half with those of the other. A Spearman–Brown correction of this half-versus-half correlation gives the statistical upper bound for a perfect model given the number of observed molecules (Methods). (G) Distribution across held-out DHSs of the Pearson correlation for mean (1D) accessibility, with 500 generated molecules per DHS. Grey, observed half versus half; red, generated versus all observed molecules; blue, Spearman– Brown upper bound. Dashed lines and numbers mark the median of each distribution. (H) As in (G), for 2D co-accessibility.

Here, we introduce a sequence-to-distribution (S2d) framework for modeling single-molecule regulatory genomics that predicts chromatin-state distributions from DNA sequence. We build a model, Mola, specifically designed for single-molecule chromatin accessibility data. We validated that Mola can accurately generate single-molecule chromatin fibers from DNA sequence and demonstrated that Mola-generated molecules capture nucleosome positioning, accessibility, DNA methylation, and distal co-accessibility between regulatory elements. We use Mola to probe genome regulatory mechanisms at the single-molecule level. By designing virtual genetic screens (e.g., motif insertions, motif combinations, and variants), we analyzed how single-molecule states relate to specific sequence features and TF grammars.

Powered by this framework, we made significant advances in these six aspects: (i) We quantified TF-motif-dependent nucleosome opening and phasing, and identify a novel nucleosome phasing TF, ZNF146, (ii) We dissected the promoter sequence grammar underlying +1 nucleosome positioning and identify YY1 as the strongest nucleosome positioning factor, (iii) We identified shared TF determinants of distal co-accessibility and co-footprinting, (iv) We uncovered a sequence-driven bimodality of chromatin states mediated by methylation-sensitive TFs, (v) We simulated pseudo-dynamic trajectories to infer the temporal scales of chromatin events, and (vi) we interpret noncoding variants beyond mean effects by analyzing distributional and distal-coupling changes. By learning distributions rather than averages of single-molecule chromatin states, this framework provides a new lens for studying the mechanisms of heterogeneous and dynamic genome regulation.

## Results

### A sequence-to-distribution framework for single-molecule regulatory genomics

Modeling single-molecule regulatory genomic data from DNA sequence is challenging because the prediction target is not a single averaged profile, but a probability distribution over all possible molecular states. This distribution describes which chromatin states can exist for a given DNA sequence, how frequently they occur, and how different states are coordinated across genomic positions. We therefore developed a sequence-to-distribution (S2d) generative modeling framework that learns to generate molecular states from this distribution (**Figure 1A**). Each single-molecule state the model generates is a sample, representing one possible single-molecule chromatin state for the input sequence. By generating many independent samples, the model creates a population of generated molecules that approximates the distribution of the ground-truth single-molecule chromatin states.

We implemented the sequence-to-distribution framework using a score-matching diffusion framework conditioned on sequence (**Methods, Figure S1**). The diffusion model is an expressive class of generative models well-suited to learning the distributions of single-molecule chromatin states^15^. During training, measured chromatin states of each molecule are progressively perturbed by noise, and the model learns to steer each noisy pattern toward chromatin states more likely for the input DNA sequence. The conditional score function defines this direction as the gradient of the log-probability density. During generation, S2d begins with random noise and iteratively applies the learned score to generate a single-molecule chromatin state. If the conditional scores are learned accurately, repeated sampling approximates the full conditional distribution (**Figure S1**). We applied this s2d framework to the thus-far deepest single-molecule regulatory genomics data in humans, the Fiber-seq dataset in human lymphocytes^16^. Fiber-seq reads have a median length of 16.6 kb, and to ensure both sufficient coverage per modeled window and long-range modeling, we used 10 kb input sequences. One can extend the input sequences into longer contexts as single-molecule read lengths increase in the future. We trained the model to take input DNA sequences and generate 10 kb single-molecule fibers highly similar to those in the training data. We name the model Mola (Model Of single-molecule Long-read chromatin Accessibility). Throughout, we refer to the single-molecule accessibility patterns within each 10 kb window as single-molecule fibers.

We demonstrated that Mola can generate accurate single-molecule fibers. To illustrate the model’s outputs, we input the 10-kb sequence around the *KIF11* promoter on the held-out chromosome (which also contains a nearby distal regulatory element) and generated 10 times as many molecules as measured in Fiber-seq for this region. Remarkably, the model-generated molecules accurately recapitulated major regulatory elements and the overall single-molecule structure (**Figure 1B**). We quantitatively evaluated Mola’s generated molecules using two complementary metrics that capture key properties of the generated distribution. First, we assessed one-dimensional (1D) accessibility, which represents the mean accessibility at each A/T position across molecules. This measure tests whether Mola accurately reproduces the average chromatin accessibility across molecules. Next, we examined two-dimensional (2D) co-accessibility, which evaluates whether Mola reproduces dependencies between positions within the same molecules. To do this, we divided each 10-kb region into bins, calculated the per-bin accessibility for each molecule, and defined co-accessibility between two bins as their Pearson correlation across molecules (**Methods, Figure S2A**). A positive correlation indicates that positions tend to be accessible together, while a negative correlation suggests mutually exclusive states. A model that predicts the correct mean but assumes independence among positions might perform well in 1D accessibility but poorly in 2D co-accessibility. While mean (accessibility) and covariance (co-accessibility) alone do not fully define a complex distribution, together they enable scalable, biologically relevant characterization of single-molecule fiber distributions. Comparing first- and second-order statistics is also a common method for assessing other types of generated data^17,18^.

Both 1D accessibility and 2D co-accessibility at the *KIF11* region were highly concordant between ground truth and model-generated fibers (1D r = 0.94, 2D r = 0.45). Notably, the 2D co-accessibility map suggests that the promoter is strongly co-accessible with the distal regulatory element, as shown by both the ground truth and the model-generated results. Increasing the number of generated molecules significantly improved the signal-to-noise ratio of co-accessibility estimates (**Figure 1C–E, S2B**). In addition to the *KIF11* promoter, we also show more random DHS regions on the held-out chromosome as examples (**Figures S3–S5**).

We further evaluated Mola’s performance on the entire held-out chromosome by generating 500 molecules per DHS. The Pearson correlation between ground truth and generated 1D accessibility was approximately 0.89, while the correlation for 2D co-accessibility was around 0.57 (**Figure 1G, H**). Because achievable correlation is limited by the number of molecules per region in the ground-truth data, especially for co-accessibility correlations, some discrepancies between generated and real fibers may reflect measurement noise and finite sampling rather than model inaccuracy. Therefore, we contextualized the correlation values by estimating their reproducibility and statistical upper bounds. To do this, we constructed a half-split benchmark by dividing the ground-truth molecules into two groups and calculating the Pearson correlation between them (**Figure 1F**). We also applied the Spearman–Brown correction to estimate a statistical upper bound (ceiling) on the correlation for both 1D accessibility and 2D co-accessibility (**Methods, Figure S2C**). Compared to this upper bound, the model achieved about 95% of the maximum performance for both metrics, indicating highly accurate predictions. In summary, we have demonstrated a novel sequence-to-distribution framework that enables prediction of single-molecule regulatory measurements for the first time. We envision this framework and the model as a powerful tool for interpreting and understanding cis-regulatory code at the single-molecule level. We demonstrate this with the following six vignettes.

### Sequence code of TF-mediated nucleosome opening and phasing

Because Mola generates single-molecule fibers only from DNA sequences, it functions as “*virtual chromatin*” enabling controlled *in silico* perturbations analogous to motif-insertion genetic screens. When only the sequence of a single motif is altered, any systematic shift in the predicted chromatin-state distribution must arise from that sequence change. This lets Mola probe cis-regulatory sequence rules with single-molecule readouts. We first quantified the impact of individual TFs on single-molecule chromatin accessibility using *in silico* motif insertion (**Figure 2A**), a common interpretability strategy for sequence-to-function models^19^. To investigate whether a de novo motif insertion can generate an accessible element, such as a DNase I hypersensitive site (DHS), and organize neighboring nucleosomes, we first selected 100 background regions outside DHSs on held-out chromosome 10 and inserted one to three copies of a TF motif at the center of each sequence. In addition, to detect insertion effects that emerge only in open chromatin regions, we selected 100 random DHS-centered backgrounds and inserted a single copy of a motif at the center of each DHS. We used 285 TF consensus motifs from a curated repertoire spanning the major human TF motif clusters^20^ (**Methods**). For each TF–background combination, we generated 100 molecules, yielding approximately 11 million molecules across diverse native sequence contexts for the following analyses.

**Figure 2.**
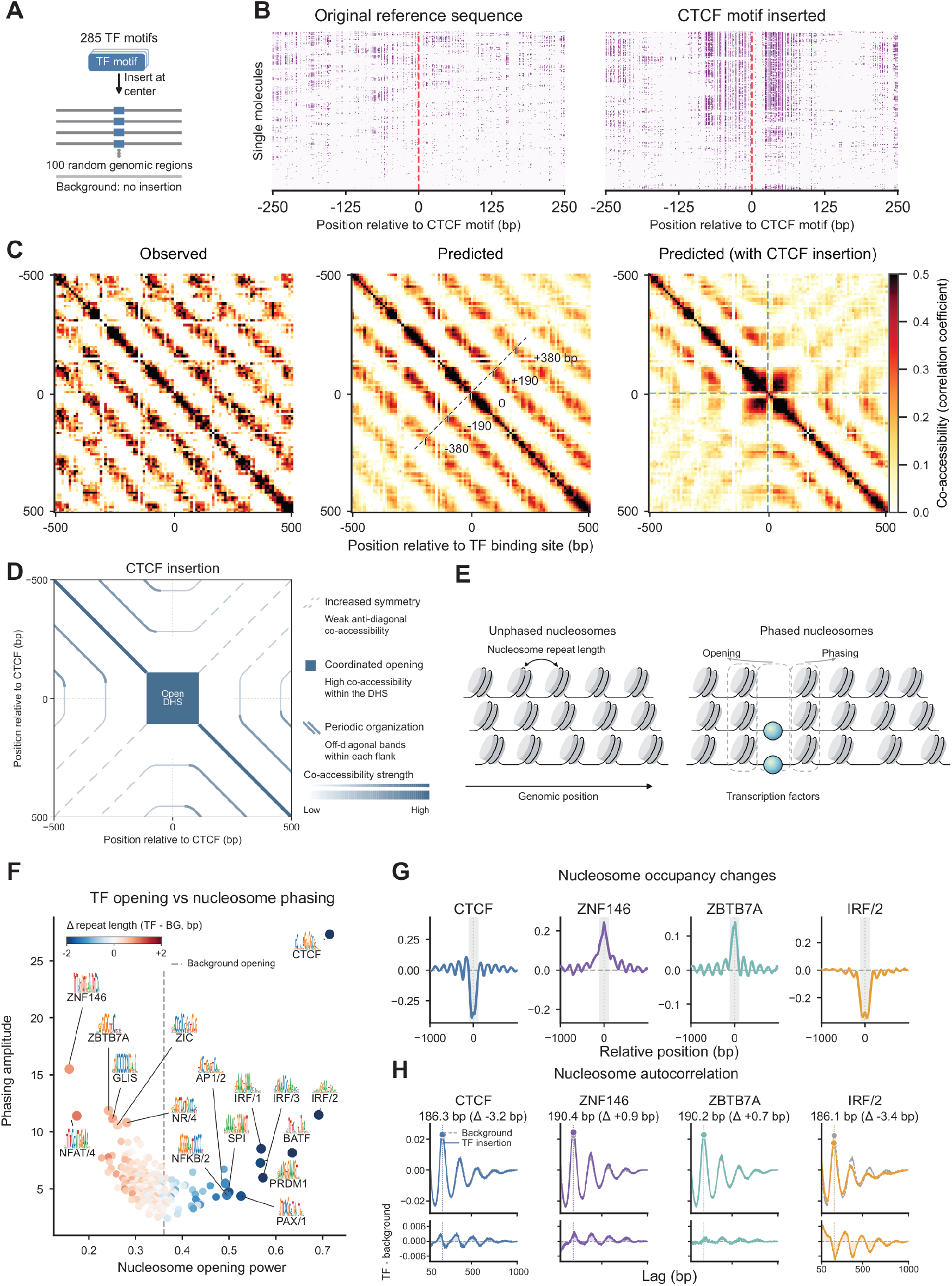
In silico motif insertion reveals how transcription factors open chromatin and organize nucleosomes. (A) Design of the motif-insertion screen. Each of 285 TF consensus motifs was inserted at the center of 100 random non-DHS background regions (one to three copies) and of 100 DHS-centered regions (one copy) on held-out chromosome 10, and Mola generated 100 molecules per sequence. (B) Generated single molecules for one background region (chr10:8,451,000–8,461,000) before (left) and after (right) insertion of one CTCF motif, shown ±250 bp around the insertion site (red dashed line); 1,000 generated molecules per panel, ordered by hierarchical clustering. Purple marks accessible positions. CTCF insertion opens the region, leaves a narrow protected footprint over the motif, and lines up the neighboring nucleosomes. (C) Co-accessibility maps (±500 bp; 10 bp bins) for the same locus, from the 56 observed Fiber-seq molecules spanning it (left), from generated molecules for the unmodified sequence (middle), and from generated molecules after insertion of one CTCF motif (right; n = 1,000 each). The dashed ruler in the middle panel marks offsets of 0, ±190, and ±380 bp from the diagonal; the stripes at these offsets come from the regular spacing of the nucleosome array. CTCF insertion creates a block of high co-accessibility at the motif and interrupts the stripes. (D) Conceptual schematic of the CTCF-insertion map in (C): a block of high co-accessibility within the open DHS (coordinated opening), periodic off-diagonal bands within each flank (nucleosome spacing), and a weaker anti-diagonal signal (increased symmetry of nucleosome organization around the motif). Line weight indicates co-accessibility strength. (E) The two effects quantified for every motif. Opening power is the gain in accessibility at the insertion site. Phasing amplitude is how regularly the flanking nucleosomes are positioned relative to the inserted motif (Methods). (F) Opening power versus phasing amplitude for one copy of each of the 285 motifs inserted into the 100 DHS-centered backgrounds (10,000 generated molecules per motif). The dashed line is the opening power of the backgrounds without insertion. Point color shows the change in nucleosome repeat length relative to background (blue, shorter; red, longer). Motif logos label the strongest openers and phasers. (G) Change in nucleosome occupancy relative to background (TF minus background) after one-copy insertion of CTCF, ZNF146, ZBTB7A, or IRF/2 into the DHS backgrounds, averaged over the 10,000 generated molecules per motif. Negative values at the motif mean nucleosome loss; positive values mean nucleosome gain. (H) Mean single-molecule autocorrelation of accessibility as a function of lag for the DHS backgrounds (grey) and after one-copy insertion of CTCF, ZNF146, ZBTB7A, or IRF/2 (color). The lag of the highest peak at lags of at least 50 bp is the nucleosome repeat length. Lower panels show the difference between the two curves (TF minus background).

Modeling chromatin accessibility at the single-molecule level allowed us to resolve TF motif effects on chromatin organization across multiple scales with high resolution. We use a CTCF motif insertion to illustrate this (**Figure 2B–E**). Before motif insertion, short tracts of accessible linker DNA were dispersed across individual molecules, revealing an organized nucleosome array with randomly positioned nucleosomes (**Figure 2B, left**). Inserting a single CTCF motif into the background sequences created a strongly accessible region centered on the insertion site, showing that the CTCF motif alone was sufficient to induce local chromatin opening in most single molecules (**Figure 2B, right**). Within this region, a narrow, protected, inaccessible genomic interval directly over the motif formed a CTCF footprint, consistent with high CTCF occupancy on open DNA. On either side of the motif, we observed strong alignment of nucleosome-sized inaccessible regions, indicating that adjacent nucleosomes occupied preferred positions relative to the motif. Thus, CTCF insertion simultaneously opened the local chromatin and established a reference point for phasing the surrounding nucleosome array, accurately recapitulating CTCF’s established chromatin-organizing activity.^21^

In addition to direct single-molecule observations, co-accessibility maps centered on TF insertion sites, calculated from single-molecule data, provide a unique view of TF effects on nucleosome organization. In background non-DHS regions, the co-accessibility maps show a clear periodicity (∼190 bp) in strong off-diagonal co-accessibility (**Figure 2C, left and middle**), indicating a regular, periodic spacing pattern of organized nucleosome arrays^22^. In contrast, CTCF insertion breaks the periodic nucleosome array into two parts separated by the open region (**Figure 2C, D**), indicating a potential physical barrier for nucleosome movement. Meanwhile, we also observe a weaker but positive co-accessibility signal along the anti-diagonal of the co-accessibility map, suggesting increased symmetry of nucleosome organization centered on CTCF binding sites (**Figure 2C, D**). In addition, we noted that CTCF insertions create high intra-DHS co-accessibility, indicating highly coordinated opening within DHS. Overall, the co-accessibility map here reflects many valuable insights into nucleosome organization around TF binding sites. We presented the impact of the top 20 TFs on co-accessibility maps as a useful resource (**Figures S6 and S7**).

For all molecules generated with different motif insertions here, we quantified two aspects of TF–nucleosome engagement: nucleosome opening and nucleosome phasing (**Methods, Figures 2E, F**). Accessibility within a cis-regulatory element reports TF-driven chromatin opening, whereas accessibility in flanking regions (mostly linker between nucleosomes) reveals varying degrees of phasing power. Interestingly, motif effects on opening and phasing appear separable, and motifs can show strong opening but weak phasing effects, and vice versa. With a single copy of motif inserted at random regions, most TFs did not strongly open chromatin or phase nucleosomes (**Figure S8 and S9A)**. However, one copy of the CTCF motif insertion was sufficient to open chromatin strongly and phase nucleosomes (**Figure 2B and S8**). These results are consistent with previous experimental results, identifying CTCF as the strongest nucleosome-opening TF^23^. We also identified cell-type-specific motifs such as IRF and BATF that opened chromatin weakly with a single copy insertion, consistent with evidence that they can act as chromatin-opening factors^24^ (**Figures S8 and S9**). The screen therefore recovered both established TF activities and factor-specific differences between opening and phasing.

The large-scale screen with motif insertions in DHSs also revealed an unexpected nucleosome-closing signature for the poorly studied factor ZNF146 (**Figure 2E, F**). Interestingly, this motif insertion can increase local nucleosome occupancy. ZNF146 was also a strong phasing factor, suggesting a distinct and unknown “nucleosome trapping + phasing” mechanism. To validate this finding, we aggregated single-molecule Fiber-seq data around ZNF146 ChIP-seq peaks and observed nucleosome-occupancy-enhancing patterns consistent with the motif insertion results (**Figure S10**), suggesting a *bona fide* role for ZNF146.

We next investigated the quantitative effects of multiple TF binding sites, since synergistic effects might reveal additional TF activity. We therefore quantified nucleosome opening and phasing for two- and three-copy motif insertions in random backgrounds. Multiple-copy insertions uncovered substantially more TFs that can open and/or phase nucleosomes, consistent with the notion of nucleosome-mediated TF cooperativity^25^. For example, three copies of IRF (specifically, we inserted the consensus sequence of the IRF/2 cluster^20^) produced substantially stronger chromatin opening than a single copy (**Figures S8 and S9**).

Opening effects by motif were broadly consistent across the two insertion backgrounds, whereas motif insertions into DHS backgrounds revealed more nucleosome-closing TFs, including the known repressive regulators ZBTB7A and ZIC^26,27^ (**Figure 2F, G, and Figure S8**). A repressor may have little measurable effect in an already nucleosome-occupied background region, but a clear effect once accessibility is established. These results suggest that ZBTB7A and related factors may fine-tune regulatory elements that are already open rather than independently create closed chromatin. In line with recent studies^28,29^, increasing nucleosome occupancy might be a universal mechanism for repressive TFs. Overall, we comprehensively quantified the impact of human TF motifs on nucleosome phasing and opening across different dosages and contexts, with implications for their distinct mechanisms.

### Quantifying TF effects on nucleosome array length and periodicity

Beyond chromatin opening and nucleosome phasing, TFs can also reshape higher-order nucleosome organization, particularly nucleosome array regularity and periodicity^30^. In human cells, nucleosomes are organized into arrays with periodic spacing of ∼180–210 bp^22^. We characterized nucleosome arrays with these two key features: nucleosome repeat length (NRL) and nucleosome periodicity strength (NPS). We quantified these features in model-generated molecules by computing the autocorrelation of the accessibility signal for each molecule (**Figure S11A**). Single-molecule data and sequence-to-distribution models are uniquely suited to analyzing the nucleosome array organization. For example, bulk-average profiles at a locus show much weaker array structure than the autocorrelation computed at the single-molecule level (**Figure S11B**).

In random non-DHS background regions, the autocorrelation of model-generated molecules yielded an estimated NRL of ∼193 bp, consistent with ground-truth data and prior reports^22^. After inserting three copies of TF motifs into background sequences, CTCF reduced NRL from ∼193 bp to ∼185 bp on average while leaving regularity relatively unaffected (**Figure S9 and S11C**). This is consistent with previous reports that depleting a CTCF-associated ATPase chromatin remodeler can significantly increase NRL length^31,32^. Notably, the chromatin-opening power of the motifs appears strongly correlated with their ability to reduce nucleosome repeat length (**Figure 2F**). In contrast to CTCF, IRF and BATF insertion reduced both nucleosome repeat length and periodicity strength, and these effects increased with motif copy number (**Figure 2H, and S11C**). Overall, effects of TF insertions on nucleosome arrays are consistent across different backgrounds, albeit weaker in DHS backgrounds. These TF-specific effects on nucleosome arrays suggest distinct mechanisms by which TFs reshape them: some primarily shift spacing while preserving regularity, whereas others both disrupt regularity and reduce spacing. In summary, molecules generated by Mola offer us a window into the versatile aspects of nucleome regulation.

### Sequence basis of +1 nucleosome positioning

Nucleosomes are positioned not only around TF-binding sites and DHSs but also around transcription start sites (TSSs). The +1 nucleosome is the first well-positioned nucleosome immediately downstream of the TSS (**Figure 3A**) and is important for promoter architecture and function^33,34^. However, the sequence basis and underlying mechanism of +1 nucleosome positioning in the human genome remain largely unclear.

**Figure 3.**
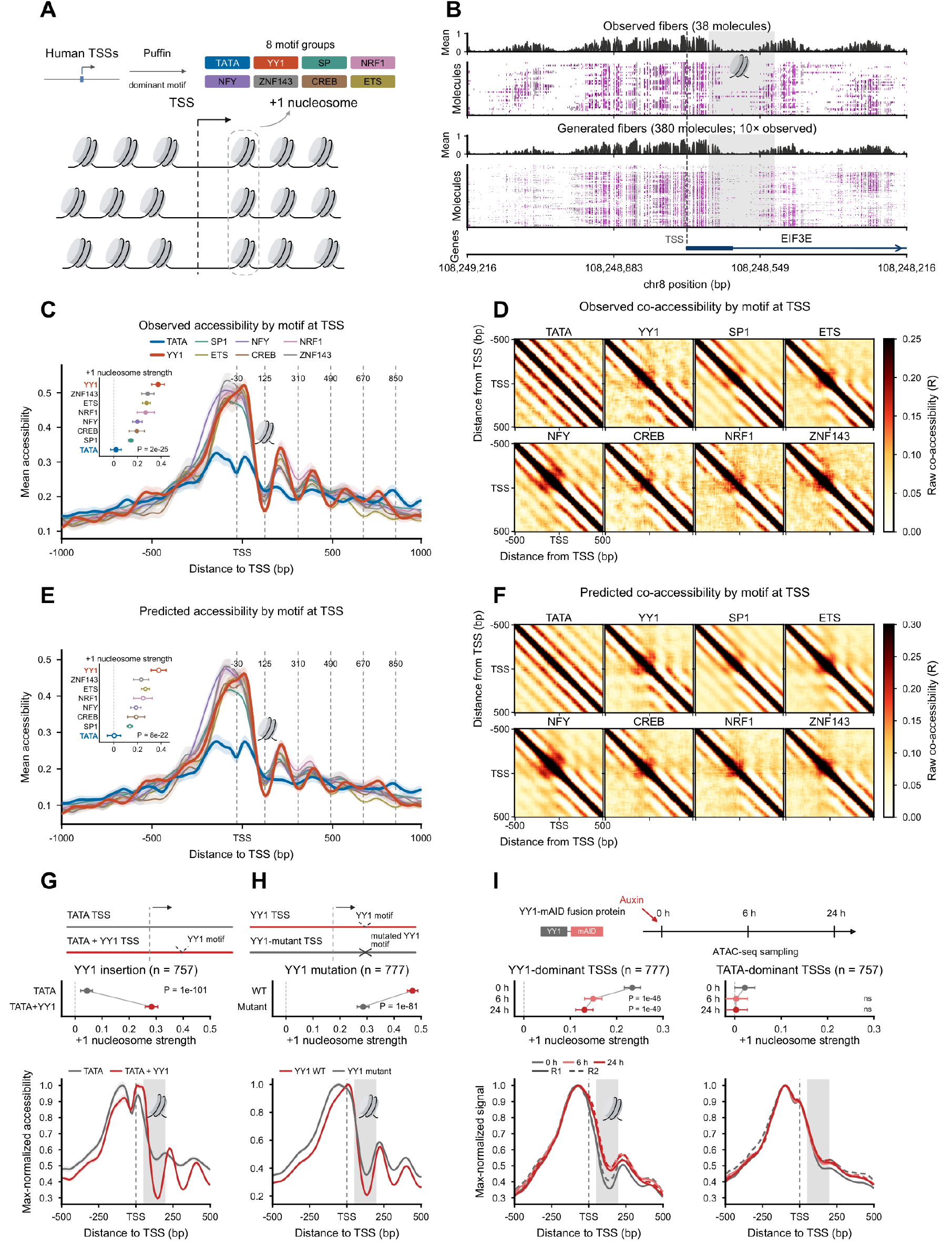
YY1 strongly determines +1 nucleosome positioning at human promoters. (A) Human TSSs were assigned to eight motif groups (TATA, YY1, SP, NRF1, NFY, ZNF143, CREB and ETS) by the dominant motif contribution to transcription initiation. Below, the +1 nucleosome is the first positioned nucleosome downstream of the TSS. (B) Observed (n = 38) and generated (n = 380) molecules at the EIF3E promoter on held-out chromosome 8. Purple, accessible positions; top tracks, mean accessibility. The gene is on the minus strand, so coordinates decrease from left to right. The dashed line marks the TSS, and the grey box marks the +1 nucleosome region (+50 to +200 bp downstream of the TSS). (C) Mean accessibility of observed molecules around the TSS (±1 kb) for each promoter group. Dashed lines mark the −30 position and the centers of the +1 to +5 nucleosomes at +125, +310, +490, +670, and +850 bp. Shading shows the standard error. The inset panel gives the +1 nucleosome strength of each group, error bars, 95% bootstrap confidence intervals over promoters, and P-values from the Kruskal–Wallis test across the eight groups. (D) Observed co-accessibility maps (±500 bp around the TSS) for each promoter group. YY1 promoters show the clearest +1 nucleosome structure; TATA promoters show almost none. (E and F) As in (C) and (D), for Mola-generated molecules (100 molecules per TSS). Inset in (E), Kruskal–Wallis P = 8 × 10⁻²². (G) In silico insertion of a YY1 motif 20 bp downstream of the TSS in all TATA-dominant promoters genome-wide (n = 757; Methods). Top, +1 nucleosome strength before (TATA, grey) and after insertion (TATA + YY1, red); error bars, 95% bootstrap confidence intervals over promoters; P, two-sided Wilcoxon signed-rank test on paired per-promoter differences. Bottom, max-normalized mean accessibility of generated molecules for the original and YY1-inserted sequences; the grey box marks the +1 nucleosome. (H) In silico mutation of the dominant YY1 motif in all YY1-dominant promoters genome-wide (n = 777). Top, +1 nucleosome strength for the wild-type (red) and mutated (grey) sequences; Bottom, max-normalized mean accessibility as in (G). (I) Analysis of ATAC-seq data from KOLF2.2J human induced pluripotent stem cells carrying a YY1-mAID degron^37^. Cells were collected 0, 6, and 24 h after auxin addition (two replicates; R1 solid, R2 dashed). Top, +1 nucleosome strength at each time point for the same YY1-dominant (n = 777) and TATA-dominant (n = 757) TSSs; error bars, 95% bootstrap confidence intervals over promoters; P, two-sided Wilcoxon signed-rank test on paired per-promoter differences versus 0 h; ns, P > 0.05. Bottom, max-normalized signal around the TSS; the grey box marks the +1 nucleosome.

We sought to ask whether Mola learns sequence determinants of +1 nucleosome positioning by generating 100 molecules for a 10 kb sequence centered on a human TSS on a held-out chromosome and piling up accessibility around the TSS. The generated molecules displayed a clear +1 nucleosome pattern consistent with ground-truth data, indicating that the model captures sequence rules underlying +1 nucleosome organization (**Figure 3B**).

To better demonstrate this and connect promoter sequence grammar to +1 positioning, we grouped TSSs into eight groups (TATA, YY1, ZNF143, SP, ETS, NFY, CREB, and NRF1) based on motif contribution score dominance from our previous transcription initiation model^35^ and compared +1 nucleosome patterns across groups (**Figure 3A, C**). Strikingly, the YY1-driven TSS (or YY1-dominated TSS) group exhibited the strongest +1 nucleosome positioning strength (**Figure 3C, E**). In contrast, TATA-driven TSSs (dominated by TBP ^36^) showed the weakest pattern, with little or weak evidence of a well-positioned +1 nucleosome, as illustrated by the *PPIF* promoter (a TATA-driven TSS) (**Figure S12**). The other six TSS groups can also position the +1 nucleosome with varying strengths, though weaker than YY1-driven TSSs. More importantly, Mola’s predicted +1 nucleosome positioning strengths correlate strongly with the ground truth across TSS groups (R = 0.99), suggesting Mola captures the sequence basis of +1 nucleosome positioning.

We also computed co-accessibility patterns around TSSs for both Mola-generated molecules and ground-truth molecules. Consistent with 1D accessibility profiles, co-accessibility maps showed the strongest +1 nucleosome-related structure at YY1-driven TSSs and nearly no such structure at TATA-driven TSSs (**Figure 3D, F**). Most TSSs exhibited intra-promoter co-accessibility within the core promoter regions, whereas TATA-driven TSSs lacked this pattern and showed lower overall accessibility despite comparable transcription initiation levels (**Figure 3D, F**). These differences were consistent between ground-truth and model-generated molecules. This suggests distinct transcription initiation mechanisms involving different modes of nucleosome organization at promoters in a sequence- and motif-dependent manner.

To further test whether the YY1 motif is sufficient to position the +1 nucleosome in the TSS, we inserted a YY1 motif into each TATA-driven TSS sequence (does not position +1 nucleosome) and generated molecules for these YY1+ TATA+ TSS sequences. The resulting fibers showed a clear, strong +1 nucleosome compared with the original TATA-only sequences (**Figure 3G**), indicating that YY1 is sufficient to position the +1 nucleosome in the promoter context. Meanwhile, we also performed *in silico* mutation of the YY1 motif sites across all YY1-driven TSS sequences and profiled the resulting molecules. We found that a YY1 motif mutation can also significantly reduce the +1 nucleosome position effects, reinforcing our conclusion here (**Figure 3H**). We further validated this model-driven conclusion with experimental data^37^. Auxin-induced degradation of YY1 in KOLF2.2J human induced pluripotent stem cells, followed by ATAC-seq, showed a strongly reduced +1 nucleosome pattern upon YY1 depletion at YY1-driven TSS but not at TATA-driven TSS, experimentally validating our conclusion here that YY1 strongly positions the +1 nucleosome (**Figure 3I**).

Finally, to explore the functional consequences of the YY1-driven +1 nucleosome, we examined transcription initiation (PRO-cap), transcriptional pausing (PRO-seq), and promoter histone marks (H3K4me3 and H3K27ac) at YY1 versus TATA TSSs. YY1 TSSs exhibited higher H3K4me3 and H3K27ac than TATA TSSs, suggesting that the +1 nucleosome might be a hotspot for promoter-related histone modifications (**Figure S13**). Meanwhile, YY1 TSSs showed lower pausing (PRO-seq) than TATA TSSs despite comparable initiation (PRO-cap), consistent with the idea that +1-nucleosome-associated chromatin features may support pause release^38^.

### Sequence basis of single-molecule co-accessibility between DHSs

Regulatory elements like DHSs operate within a broader sequence context, often comprising multiple promoters and enhancers that may act together at the single-molecule level. While bulk accessibility assays measure the average activity of each element, they do not reveal whether two elements tend to open simultaneously on the same molecules. Long-read single-molecule techniques such as Fiber-seq allow direct assessment of this relationship through co-accessibility, which is the correlation in accessibility between two regions along individual chromatin fibers. Overall, within the 10 kb window, co-accessibility for all pairs of DHSs showed little distance dependence (**Figure S14**), consistent with model predictions. However, the sequence basis that determines co-accessibility among regulatory elements (i.e., DHSs) remains unclear. To address this, Mola generates extensive single-molecule data from perturbed sequences, thereby providing a scalable approach for identifying sequence features that encode co-accessibility. We examined whether the coordination between two DHSs depends on the types and combinations of their TF motifs. Specifically, we selected 19 TFs with the strongest effects on local accessibility. We inserted their motifs at two sites within 100 randomly selected background sequences, creating two synthetic DHSs. The distance between two sites ranged from 1 kb to 8 kb (**Figure 4A**). After generating fibers with these synthetic sequences by Mola, we assessed co-accessibility between the two DHSs across molecules. Since each synthetic DHS was defined by a specific TF motif, we refer to it as a TF-DHS.

**Figure 4.**
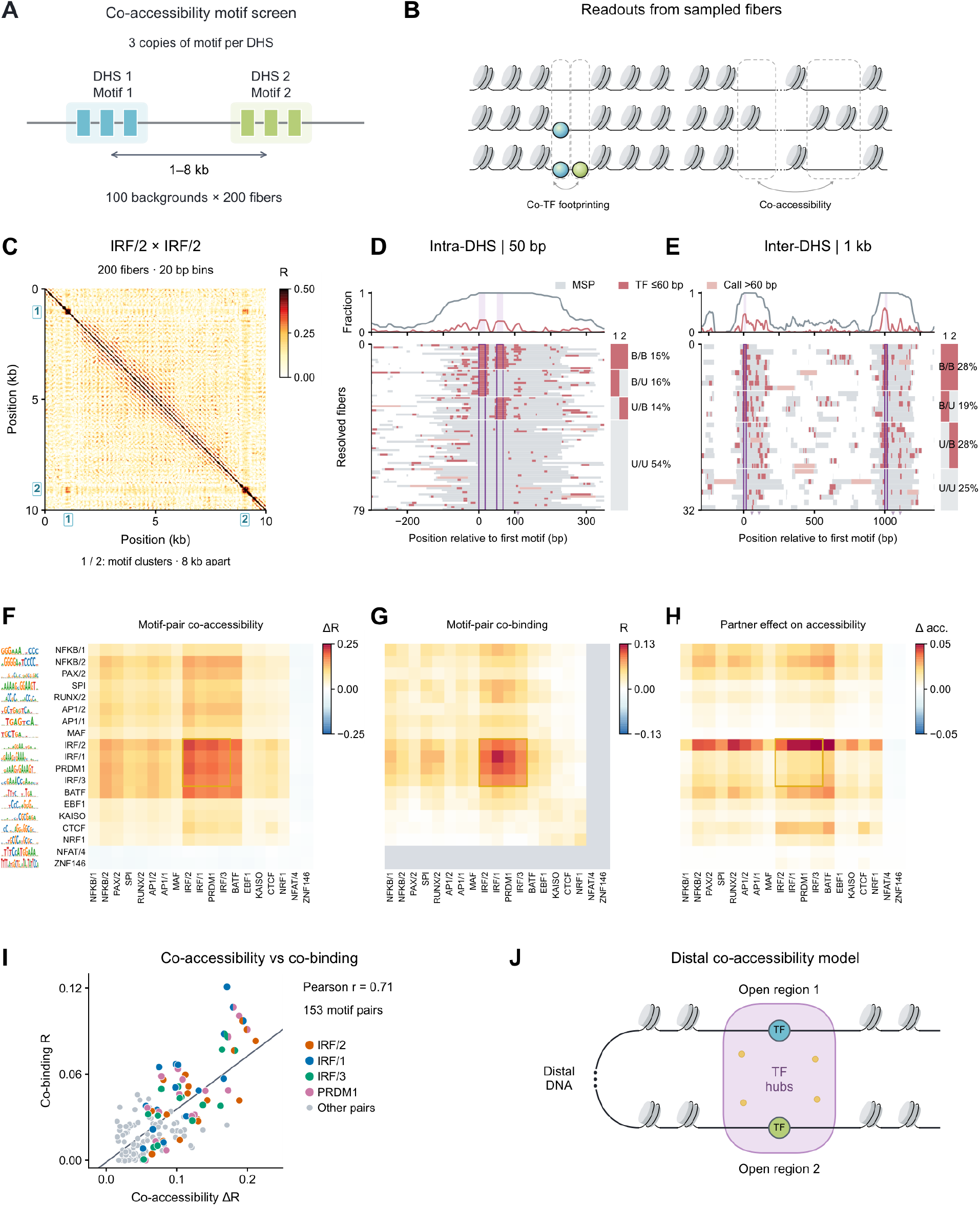
Shared sequence basis of single-molecule co-accessibility and co-binding between distal DHSs. (A) Design of the two-DHS screen. For every pair of 19 TF motifs, three copies of motif 1 and three copies of motif 2 were inserted 1–8 kb apart into 100 background sequences to create two synthetic DHSs, and 200 molecules were generated per sequence. (B) Two readouts taken from the generated molecules. Co-binding asks whether both motifs carry a TF footprint on the same molecule. Co-accessibility asks whether both DHSs are open on the same molecule. (C) Co-accessibility map (20 bp bins; 200 molecules) for a sequence carrying two IRF/2 motif clusters 8 kb apart (positions 1 and 2). The off-diagonal signal in the corners shows that the two synthetic DHSs open together. (D) FiberHMM footprint calls on generated molecules for one background region carrying two IRF/2 motifs 50 bp apart within the same DHS. Each row is a molecule. Grey, methyltransferase-sensitive patches (accessible DNA); red, TF footprints of at most 60 bp; purple bars, motif positions. Rows are grouped by the binding state of motifs 1 and 2 (B, bound; U, unbound): B/B 15%, B/U 16%, U/B 14% and U/U 54% of testable molecules. The top track gives the fraction of molecules with each feature at each position. (E) As in (D), for two IRF/2 motifs in separate DHSs 1 kb apart (B/B 28%, B/U 19%, U/B 28%, U/U 25%). Light red marks protected segments longer than 60 bp. (F) Motif-pair co-accessibility matrix for the 19 TFs. Each cell is the mean co-accessibility between the two synthetic DHSs (ΔR, change in Pearson correlation relative to background), averaged over backgrounds and distances. Motif logos and row labels on the left apply to (F–H). The IRF-family block is highlighted. (G) Motif-pair co-binding matrix. Each cell is the correlation of footprint occupancy between the two motifs across molecules. NFAT/4 and ZNF146 (grey) were excluded because they close chromatin. (H) Partner-induced accessibility change. Each cell is the change in mean accessibility of the target DHS (column) when a partner DHS (row) is inserted 1–8 kb away, relative to the target inserted alone, averaged over distances and backgrounds. Positive values mean the partner makes the target more accessible. (I) Co-binding versus co-accessibility for the 153 TF pairs in the matched 17 × 17 matrices (Pearson r = 0.71). Pairs that include IRF/1, IRF/2, IRF/3, or PRDM1 are colored. (J) Model. Compatible TFs bound at two distal DHSs may be held together in a shared TF hub, a region of high local TF concentration.

To identify TF motifs that preferentially drive coordinated opening between distal regions, we constructed a TF-by-TF co-accessibility strength matrix based on the Mola-generated molecules (**Figure 4F**). Each element of the matrix represents the average co-accessibility strength between the row TF-DHS and the column TF-DHS across different background regions and genomic distances. This TF-by-TF co-accessibility matrix revealed TF-specific contributions to forming co-accessibility. For example, IRF family factors (i.e., IRF/1, IRF/2) motif-based DHS pairs exhibit the highest co-accessibility with each other. In contrast, although CTCF had the strongest DHS accessibility (**Figure 2**), it did not yield the strongest co-accessibility pattern, indicating that coordinated opening is not simply an additive or multiplicative product of local accessibility. Repressive TFs like ZNF146 do not promote co-accessibility with any other TFs tested here.

We also validated this TF-by-TF co-accessibility strength matrix by examining and quantifying the co-accessibility of all the possible pairs of TF-TF DHS sites within 10 kb in the ground-truth Fiber-seq data (**Methods, Figure S15**). For example, the model-generated co-accessibility maps of CTCF DHSs closely recapitulated pile-up co-accessibility maps derived from ground-truth Fiber-seq molecules centered on CTCF motif positions (**Figure S16**). Notably, both generated and observed CTCF co-accessibility maps exhibited a regular grid-like pattern, consistent with the recent report of fiber homotypy, a phenomenon of long-range coordination of nucleosome spacing patterns^39^. Importantly, the Fiber-seq-based TF-by-TF co-accessibility matrix is highly concordant with Mola’s results (r = 0.7). These results support a model in which TF identity and combinatorial grammar strongly influence molecule-wise coordination between distal elements.

We then investigated the functional impact of co-accessibility between DHSs, specifically whether a highly co-accessible partner DHS generally enhances local accessibility at the target DHS. To assess this, we compared mean accessibility of the target DHSs across various partner-DHS insertions. Our results showed that for most TF-DHSs, inserting a partner distal DHS or altering its identity only minimally affected the mean accessibility at the original anchor site (**Figure 4H, S16**). However, for motifs we found to strongly promote co-accessibility, such as IRF, inserting the second DHS at one anchor site often substantially increased accessibility at the other anchor. This suggests that coordinated TF activity can stabilize or enhance the accessibility of a partner DHS. These findings highlight the important role of TF identity in determining distal coordination between regulatory elements.

### Shared sequence basis of TF-driven co-binding and co-accessibility

The TF-specific co-accessibility patterns raised the hypothesis that coordinated opening is accompanied by coordinated occupancy of the underlying TFs. We therefore asked whether motif pairs that produced strong co-accessibility also exhibited preferential co-binding (or co-footprinting) on the same generated molecules (**Figure 4B**). IRF served as a representative example, as homotypic IRF insertion produced the strongest co-accessibility signal in our screen (**Figure 4C**). To analyze the binding states of each inserted motif on individual molecules, we employed FiberHMM^40^ (**Methods; Figure S17**). When two IRF/2 motifs were separated by 50 bp within the same DHS, 347 molecules (pooled across the 100 backgrounds) exhibited footprints at both sites, compared with 262.6 expected from their marginal binding frequencies (32.1% increase; P = 1.39 × 10⁻¹⁰; **Figure 4D**). When the motifs were in separate DHSs 1 kb apart, 297 molecules were bound at both sites, compared with 238.5 expected under independent binding (24.5% increase; P = 1.81 × 10⁻⁶; **Figure 4E**). Generally, we observed that inter-DHS co-binding between TFs is slightly weaker than intra-DHS co-occupancy (**Figure S18**). These findings suggest that IRF motifs not only facilitate chromatin opening but also promote TF co-occupancy across distal regulatory elements, indicating a shared long-distance coordination mechanism.

We extended the analysis across different TF identities by creating a TF-by-TF co-binding matrix for each single-molecule fiber generated by Mola. Each element of this matrix indicates the correlation between binding events at two TF-DHSs (**Methods**). Complete co-binding data were available for 17 of the 19 TF families included in the co-accessibility matrix; NFAT (NFAT/4) and ZNF146 were excluded from the co-binding matrix because they repress open chromatin (**Figure 4F, G**). When comparing the co-binding matrix to the co-accessibility matrix, co-binding and co-accessibility were strongly concordant across the 153 TF-pair comparisons in the matched 17 × 17 matrices (r = 0.71; **Figure 4I**). Notably, IRF-family motifs (IRF/1, IRF/2, IRF/3, and PRDM1) showed both strong co-accessibility and significant co-binding. The co-binding patterns of these TF pairs in the ground-truth Fiber-seq data also aligned closely with Mola’s results (**Figure S18**).

This concordance supports a shared sequence-encoded basis for coordinated TF occupancy and coordinated chromatin opening. These observations may be mediated by TF-regulatory hubs that act over long distances (> 10 kb), potentially through high TF concentration-mediated local assemblies^41^, that stabilize TF occupancy and promote distal DHSs in an accessible state on the same molecule (**Figure 4J**). This model provides a potential mechanism through which sequence-specific TF interactions could support distal coordination of regulatory elements, although the biophysical and biochemical nature of such hubs remains to be tested.

### Single-molecule joint modeling of DNA methylation and accessibility

Chromatin accessibility and DNA (CpG) methylation are well known to be inversely correlated at the cell-population average level. However, their relationship at the single-molecule level at the same locus has been much less studied^42^. Because Fiber-seq can measure exogenous m6A accessibility and endogenous m5C methylation on the same chromatin fibers, we extended Mola to model their joint distribution at single-molecule resolution. This allowed us to ask not only whether Mola predicts each modality separately, but also whether DNA sequence predicts how accessibility and methylation are coupled within individual regulatory elements.

We trained Mola to predict methylation and accessibility jointly, so each generated molecule consists of an accessibility track and a CpG methylation track that can be directly compared with observed molecules. For example, on a held-out region, generated molecules faithfully reproduced the locus’s observed accessibility and methylation patterns at the single-molecule level (**Figure 5A**), as well as the relationship between accessibility and methylation across molecules (**Figure 5B**). Across 10,029 held-out regions on a genome-wide scale, the generated mean methylation profile matched the observed profile (mean r = 0.76, 79% of the upper bound; **Figure S19**). Most importantly, the model captured how the two modalities relate within each element. The cross-molecule correlation between accessibility and methylation computed from generated molecules agreed with that computed from observed molecules (r = 0.38, 69% of the upper bound; **Figure 5C**). Mola therefore predicts not only each modality but also their coupling on the same molecule from sequence alone.

**Figure 5.**
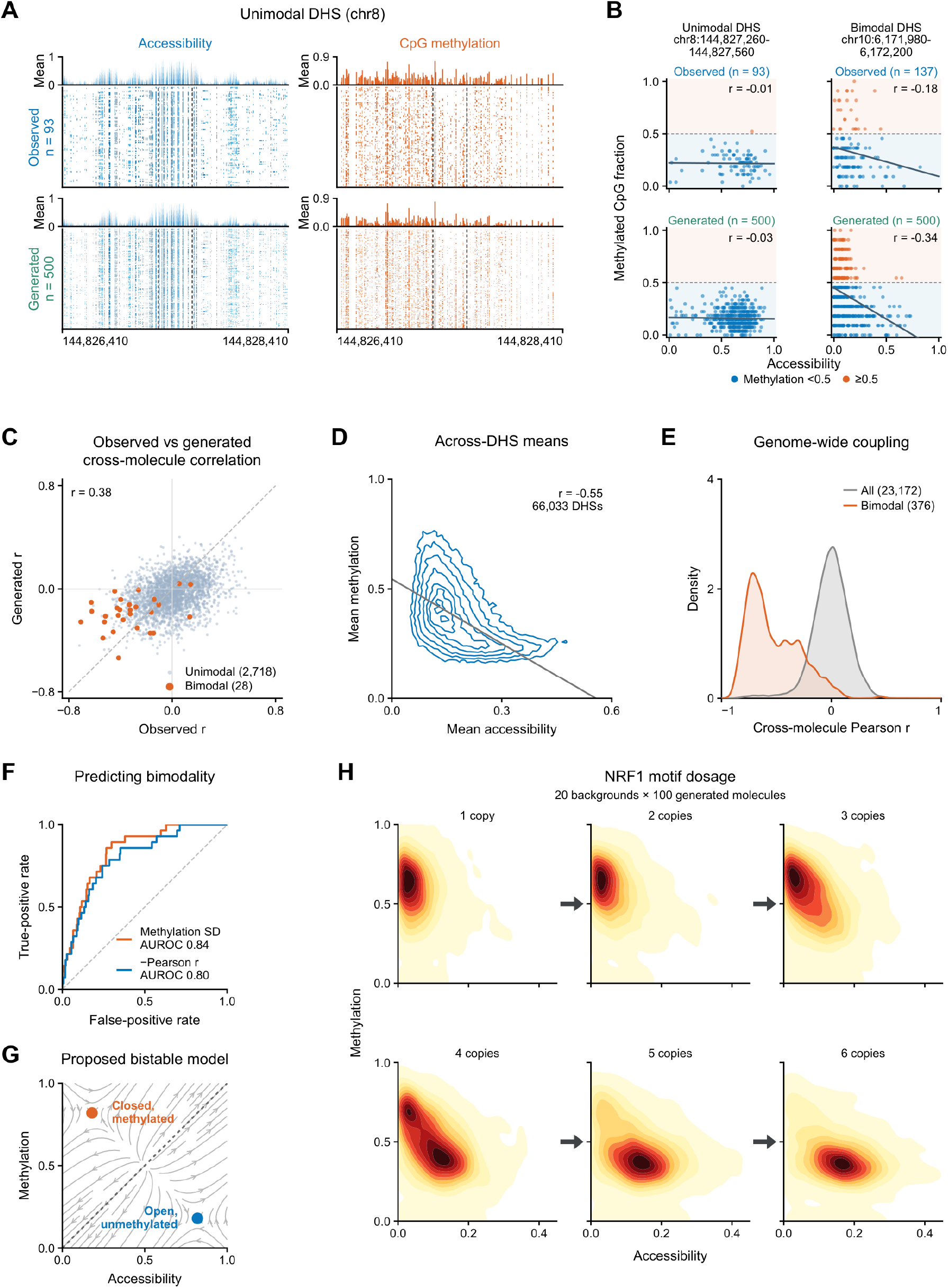
Joint modeling of single-molecule chromatin accessibility and CpG methylation. (A) Observed (n = 93) and generated (n = 500) molecules at a unimodal DHS on held-out chromosome 8 (window chr8:144,826,410– 144,828,410; DHS chr8:144,827,260–144,827,560, dashed lines). Left, accessibility (blue); right, CpG methylation (orange) on the same molecules. Mean tracks are shown above each molecule panel. (B) Per-molecule accessibility versus methylated-CpG fraction at the unimodal DHS shown in (A) (left) and at a bimodal DHS on chromosome 10 (chr10:6,171,980–6,172,200; right), for observed (top) and generated (bottom) molecules. Points are colored by methylation below (blue) or above (orange) 0.5; lines are linear fits; r, Pearson correlation. (C) Cross-molecule accessibility–methylation correlation computed from generated molecules versus the same correlation computed from observed molecules, for 2,746 held-out DHSs (correlation across DHSs, r = 0.38). Orange, the 28 DHSs called bimodal from observed molecules; (D) Mean methylation versus mean accessibility in observed Fiber-seq molecules across 66,033 DHS-centered 1 kb windows genome-wide (density contours; r = −0.55). (E) Distribution of cross-molecule correlations in observed molecules for all 23,172 testable DHSs (grey) and for the 376 bimodal DHSs (orange). (F) ROC curves for predicting which held-out DHSs are bimodal (28 bimodal versus 2,702 unimodal DHSs) from two scores computed on generated molecules only: the standard deviation of methylation across molecules (AUROC 0.84) and the negative cross-molecule correlation (AUROC 0.80). (G) Proposed bistable model. A closed, methylated state and an open, unmethylated state are separated by an unstable boundary; arrows show the direction of change from different starting points. Methylation blocks NRF1 binding, and NRF1 binding promotes accessibility and loss of methylation, so each state reinforces itself. (H) Joint distribution of per-molecule accessibility and methylation within 600 bp of the insertion site after inserting one to six copies of the NRF1 motif into 20 background regions (100 generated molecules each). One or two copies leave molecules closed and methylated; three or four copies give a bimodal distribution; five or six copies move molecules to the open, unmethylated state.

We next examined the coupling between methylation and accessibility more carefully. Across genome-wide DHSs, mean accessibility and mean methylation over one DHS were anticorrelated as expected (Pearson r = −0.55; **Figure 5D**). Across molecules at one DHS, however, accessibility and methylation varied almost independently from molecule to molecule. The median correlation coefficient across molecules was zero (**Figure 5E**). Interestingly, a minority of DHSs still showed significant negative correlation between methylation and accessibility across molecules (**Figure 5B**).

We speculate that this minor group of negatively correlated DHSs has two states of DNA methylation (**Figure 5B**). To better analyze these bimodal regions, we quantified the bimodality of each DHS based on per-molecule methylation profiles (**Methods**). Genome-wide, 376 of 23,172 testable DHSs (1.6%) were two-state (bimodal). Nearly all of them were anticorrelated; 372 (98.9%) had a negative cross-molecule correlation, with a median of −0.55 compared with −0.05 at unimodal DHSs (**Figure 5E**). Molecule-level coupling between accessibility and methylation is thus concentrated in two-state DHSs, where each molecule is either closed and methylated or open and unmethylated. The two example loci in Figure 5B illustrate the contrast: at a two-state DHS, observed molecules split into two methylation states, and the less methylated molecules were more accessible and generated molecules reproduced this coupling.

We next asked whether this bimodality can be predicted from sequence. The generated molecules by Mola for the 28 identified bimodal DHSs on the held-out chromosomes also showed strong anticorrelation and greater methylation variation, consistent with the observed molecules (26 of 28 are negatively correlated; median r = −0.18 versus −0.03 at unimodal DHSs; **Figures 5C and S20**). Consequently, two scores computed only from generated molecules, the variability (s.d.) of methylation across molecules and the strength of its anticorrelation, can predict the observed two-state DHSs (AUROC 0.84 and 0.80; **Figure 5F**). These results show that Mola can predict DHS bimodality from sequences.

To further identify sequence determinants of the bimodality, we first performed motif enrichment analyses between bimodal DHSs and unimodal controls. NRF1 and ZBTB14 were the most significantly enriched motifs (**Figure S21**). In addition, Mola-based motif screening by inserting motifs into random genomic backgrounds showed that NRF1 insertion produced the strongest accessibility–methylation anticorrelation among 285 tested TF motifs (r = −0.44), followed by KAISO (r = −0.43; **Figure S22**).

We thus propose a potential mechanism for sequence-based bimodality: a bistable epigenetic landscape created by methylation-sensitive TF motifs. The two stable states include a closed, highly methylated state and an open, hypomethylated state (**Figure 5G**). Given the strong methylation sensitivity of NRF1^43^, CpG methylation would reduce NRF1 binding, whereas NRF1 occupancy would promote accessibility and loss of methylation, thereby pushing the single-molecule toward the two stable states. Feedback among TF occupancy, accessibility, and methylation maintenance therefore stabilizes both states.

This mechanism predicted that the bimodality would be motif dosage-sensitive. Indeed, in model-based motif-insertion screening, we found that increasing NRF1 motif dosage from 1 to 6 progressively reshaped the joint accessibility-methylation distribution (**Figure 5H**). With one or two motifs, molecules remained concentrated in a relatively inaccessible, highly methylated state. Intermediate copy numbers broadened the distribution into a bimodal shape along the antagonistic axis, whereas five- or six-motif insertions shifted the molecules toward high accessibility and low methylation (**Figure 5H**). Motif dosage can therefore alter the joint distribution of single-molecule accessibility and methylation states. Together, these results show that Mola learns the sequence basis of bimodal accessibility-methylation states.

### Simulating dynamic trajectories with a sequence-to-distribution model

Single-molecule sequencing provides static snapshots of chromatin states, but does not directly measure the temporal order of regulatory events, such as TF binding and nucleosome movement along DNA. Although static snapshots cannot reveal the true timing of biological events, the learned distribution captures which chromatin configurations are probable and how probability changes between neighboring states. The model’s score function uses this information to guide each state toward nearby, more probable states. Langevin dynamics can combine this guidance with random fluctuations to move through the learned state space, producing statistically plausible trajectories that will converge to the learned distribution rather than arbitrary orderings of static samples. To connect the model-learned distribution to pseudo-dynamic trajectories, we used the model’s learned score function to guide Markov chain Monte Carlo sampling with Langevin dynamics (**Figure 6A**). Briefly, starting from an initial sample generated by Mola, we updated each state through a combination of score-directed drift toward high-probability chromatin states and stochastic noise, enabling transitions between states and exploration of the broader distribution. The resulting ordered samples therefore trace plausible transition routes through the model-implied chromatin state space, providing pseudo-dynamic-based trajectories.

**Figure 6.**
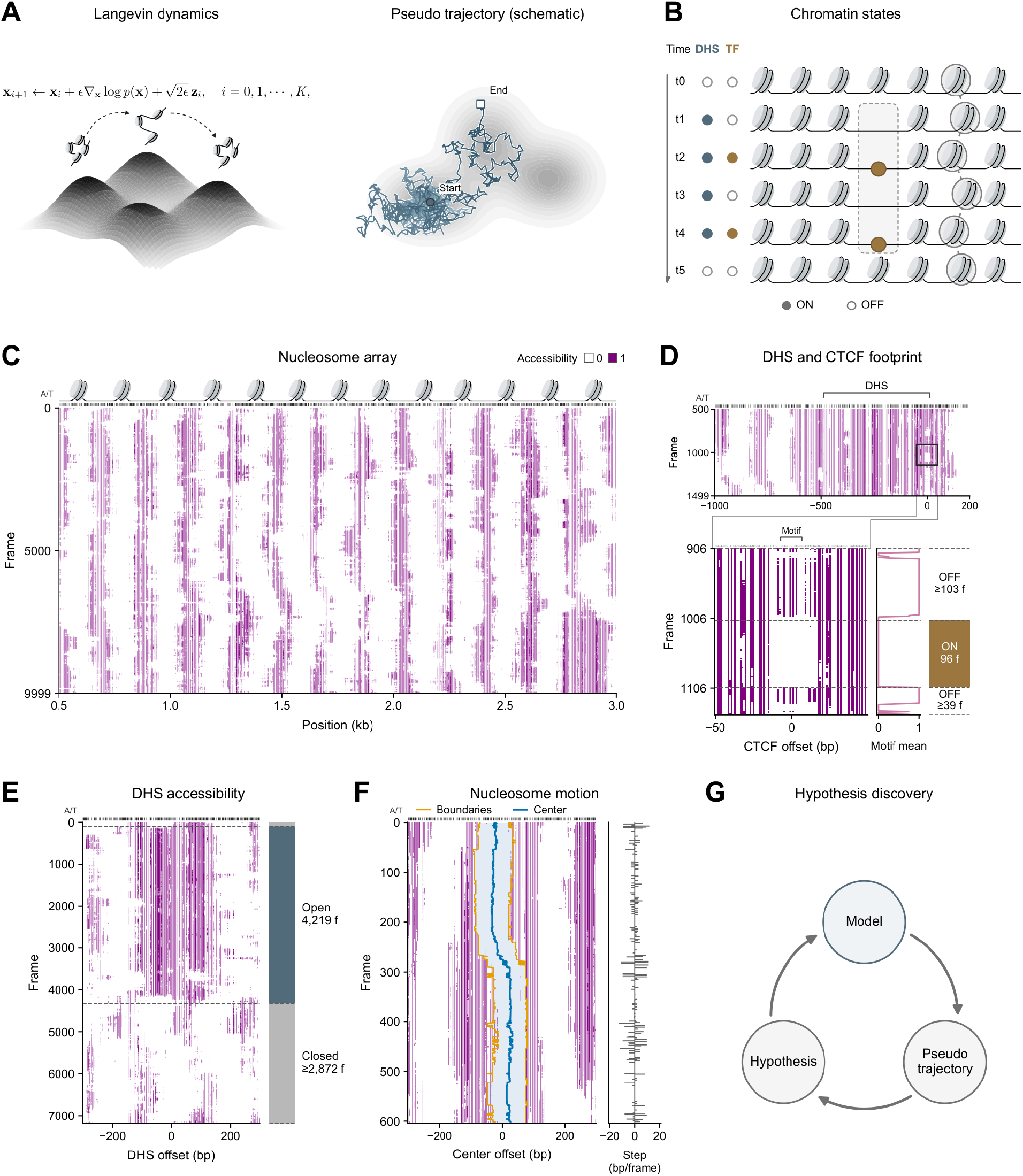
Pseudo-dynamic trajectories sampled from the learned distribution. (A) Left, Langevin sampling. Starting from a generated molecule, each update moves the chromatin state along the model score (the gradient of the log probability) and adds random noise, so the chain travels between probable states. Right, a schematic trajectory in an illustrative two-dimensional density. (B) Events that can be read from a trajectory: DHS opening and closing, TF footprint appearance and disappearance within an open DHS, and nucleosome movement along the DNA. (C) Nucleosome array along one trajectory in the NAPA region (chr19:47,509,500–47,512,000); all trajectories in (C–F) were sampled for the same 10-kb window (chr19:47,509,000–47,519,000). Rows are 10,000 consecutive frames and columns are genomic positions; purple, accessible; white, inaccessible (rounded model output, no smoothing). Icons above mark the dominant inferred nucleosome positions over the whole trajectory, and the strip marks reference A/T positions. One frame is one Langevin update. (D) DHS and CTCF footprint along a second trajectory. Top, frames 500–1,499 across the DHS (chr19:47,515,100–47,515,600), with positions given relative to the CTCF motif center. The boxed 100-bp window and frame interval are expanded below (frames 906– 1,106), with the motif position marked. To the right of the zoom, the mean accessibility over the motif’s A/T positions and the footprint state, called ON (ochre) or OFF (pink), in each frame, with run lengths in frames (f). (E) DHS accessibility along a third trajectory, ±300 bp around a 193-bp DHS (chr19:47,518,230–47,518,423). The strip calls each frame Open (blue) or Closed (grey) from FiberHMM nucleosome coverage of the DHS (Methods), with run lengths in frames. (F) Nucleosome motion along a fourth trajectory. Left, accessibility around one nucleosome over 600 frames, with the inferred footprint (shaded), its boundaries (orange), and its center (blue). Right, frame-to-frame change in center position. This footprint’s center shifted 62.5 bp over 100 frames. (G) Proposed discovery cycle.

One can analyze three key aspects of chromatin events from these pseudo-dynamic trajectories. At each DHS site, these pseudo-trajectories capture the dynamics of chromatin accessibility, including DHS opening and closing. Within an open DHS site, the pseudo-trajectories also capture TF-footprint appearance and disappearance, suggesting TF-binding kinetics. Beyond DHS regions, pseudo-dynamic trajectories also reveal nucleosome diffusion/movement along DNA (**Figure 6B**). To illustrate our framework, we generated 100 pseudo-trajectories in the *NAPA* promoter region that contain both a strong CTCF site and intermediate-level DHS. Visualizing these trajectories clearly showed the on-and-off dynamics of a CTCF footprint and nucleosome movement along the DNA across the simulated trajectory (**Figure 6C, D**).

We next quantified these three key features with the pseudo-dynamic trajectories. At the CTCF motif, footprint occupancy switched rapidly, with an estimated median footprint duration of 153 (95% confidence interval, 127–174) and median no-footprint duration of 151 frames (95% CI: 130–187), respectively (**Figure 6D**). By contrast, at the DHS level, DHS opening/closing switched much more slowly (**Figure 6E**), with the estimated median duration of 3,186 frames for open states (95% confidence interval, 1,929–5,219) and 1,876 frames for closed states (1,380– 3,627). The longer DHS accessibility duration relative to TF binding is consistent with previous chromatin-remodeler inhibition studies showing that changes in DHS accessibility occur over tens of minutes to several hours^44,45^.

Beyond the DHS region, we also tracked the nucleosome footprints to characterize nucleosome movement (i.e., diffusion) relative to DNA sequence position (**Figure 6F**). For example, a relatively mobile nucleosome footprint shifted by 62.5 bp over 100 frames. Across all tracked nucleosome trajectories, positional changes were generally modest. At a lag of 100 frames, the mean absolute displacement was 8.2 bp, and the root-mean-square displacement was 11.7 bp; because this metric weights large movements more heavily, it indicates that most nucleosomes moved only modestly while a smaller subset underwent substantially larger displacements. To examine how movement accumulated, we calculated mean-squared displacement over increasing frame intervals. Unlike ordinary one-dimensional Brownian diffusion, for which mean-squared displacement (MSD) increases linearly with time, the nucleosome MSD increased sublinearly, with a fitted scaling exponent of approximately 0.57 over lags up to 200 frames. We interpret this pattern as evidence of constrained, subdiffusive nucleosome repositioning along DNA. This nucleosome pattern along these pseudo-trajectories is consistent with optical-tweezer experiments demonstrating confined base-pair-scale nucleosome diffusion along DNA^46^.

Together, these results reveal a temporal scale hierarchy of chromatin pseudo-dynamics that the model captures. CTCF footprints appear and disappear rapidly within DHSs that remain open or closed for much longer periods, while nucleosomes generally move over short distances with occasional larger shifts. They provide testable predictions about the relative time-scale of chromatin dynamic events. To the best of our knowledge, no experimental method can probe these chromatin dynamics *in situ,* let alone genome-wide. However, this framework of model-based pseudo-trajectories provides a roadmap toward this goal through an iterative, model-guided discovery cycle. In this cycle, a dynamic model generates pseudo-trajectories consistent with the observed distribution, yielding measurable and falsifiable predictions that, in turn, motivate new analyses or experiments; these new experimental data further refine the mechanistic hypothesis and improve the dynamic modeling approaches (**Figure 6G**).

### Dissecting genetic variant impacts at the single-molecule level

A key application of sequence models is large-scale predictions and interpretations of noncoding variants^6,8^. Existing sequence-to-function models predict only the mean effects of a variant and therefore cannot reveal whether a variant changes chromatin-state distribution or coordination between regulatory elements on the same molecule. Mola overcomes this limitation by generating single-molecule chromatin states for reference and alternative alleles, thereby extending quantification of variant effects from mean accessibility to higher-order effects, including co-accessibility and nucleosome array organization (**Figure 7A**).

**Figure 7.**
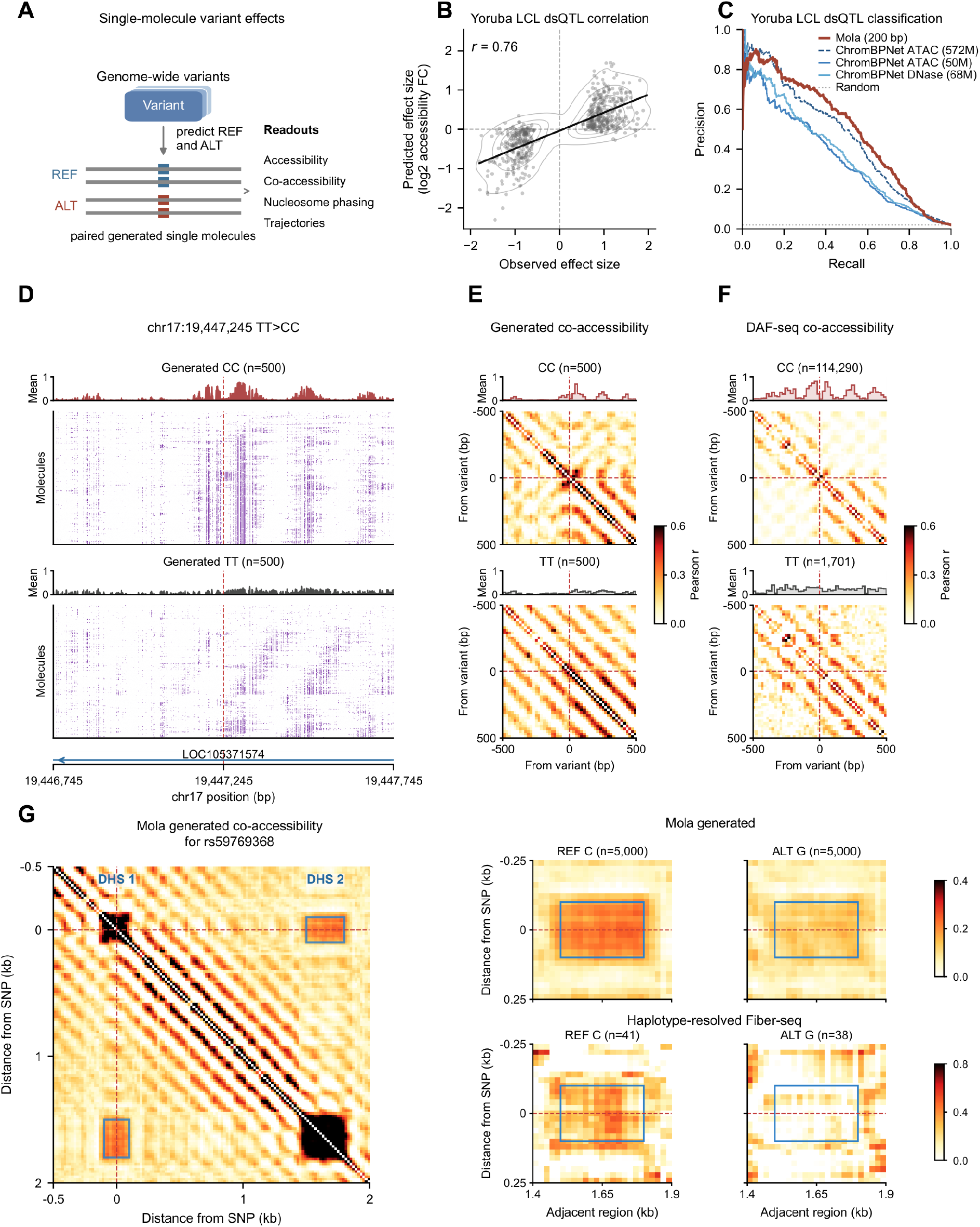
Mola predicts variant effects on single-molecule accessibility and distal co-accessibility. (A) For each variant, Mola generates matched populations of molecules for the reference (REF) and alternative (ALT) alleles. Effects are read out as differences in mean accessibility, co-accessibility, nucleosome phasing, and trajectories. (B) Predicted versus observed effect size (log2 fold change in accessibility) for 573 Yoruba LCL dsQTLs (Pearson r = 0.76). Line, linear fit; contours, point density. (C) Precision–recall curves for classifying 27,373 variants as dsQTLs using the absolute predicted effect. Mola (red) is compared with ChromBPNet models trained on ATAC-seq (572M and 50M reads) and DNase-seq (68M reads). The dotted line is the random baseline (0.020). (D) A melanoma-associated somatic CC-to-TT mutation at chr17:19,447,245–19,447,246 within a CTCF motif. Generated molecules (n = 500 per allele) for the reference CC and mutant TT alleles, ±500 bp around the variant (red dashed line). Purple, accessible positions. (E) Co-accessibility maps computed from the generated molecules for each allele. (F) Co-accessibility maps computed from published DAF-seq single-molecule footprinting at the same locus (reference CC, n = 114,290 molecules; mutant TT, n = 1,701). In both prediction and data, the TT allele loses the CTCF-centered pattern and gains co-accessibility across the site. (G) The caQTL rs59769368. Left, Mola co-accessibility map (−0.5 to 2 kb relative to the variant) for the reference allele, showing the variant-containing DHS 1 and a second DHS about 1.5–1.8 kb downstream. Blue boxes mark the DHS 1–DHS 2 interaction. Right, zoom in on this interaction for the reference C and alternative G alleles in Mola-generated molecules (top; n = 5,000 per allele) and in haplotype-resolved GM12878 Fiber-seq molecules (bottom; n = 41 and 38). Co-accessibility between the two DHSs is much weaker on the G allele in both.

To evaluate Mola’s capacity for variant effect prediction, we first showed that Mola accurately predicts conventional mean-accessibility effects, comparable to ChromBPNet^47^. Across 573 Yoruba LCL dsQTLs^48^, Mola predicted reference-to-alternative log2 accessibility fold changes were strongly concordant with experimentally measured effects (r = 0.76, **Figure 7B**), on par with the performance of expert models in mean accessibility (ChromBPNet) (**Figure S23**). In a larger classification benchmark comprising 27,373 variants in the same study, the absolute Mola effect score achieved an average precision of 0.493. This exceeded the average precision of mean predictions from ChromBPNet (0.353–0.462) (**Figure 7C**). Predicted effects were also concordant across 6,821 African LCL caQTLs^49^(r = 0.67, **Figure S23**), establishing that Mola performs competitively on the average variant-effect tasks addressed by existing sequence models.

Mola’s key advance is its ability to predict how variants reshape single-molecule chromatin accessibility, particularly co-accessibility between individual molecules, which no model thus far can predict and for which no genome-scale experimental benchmark currently exists. We first demonstrated this with a melanoma-associated somatic mutation, previously profiled with ultra-high-depth single-molecule footprinting^50^. Mola’s predictions for both single-molecule and co-accessibility are highly consistent with the experiment (**Figure 7D–F**). Mola accurately predicted the decrease in mean accessibility profile for the TT allele relative to the CC allele, consistent with the TT variant disrupting the CTCF motif locally (**Figure 7D**). Importantly, Mola predicted increased co-accessibility between the two sides and a loss of the antidiagonal co-accessibility pattern expected with CTCF disruption (**Figure 7E**, **Figure 2C**). Mola therefore precisely predicts the variant’s role in reorganizing the nucleosome array across individual molecules.

To further evaluate Mola’s capability in predicting co-accessibility between DHSs, we searched across known dsQTLs and caQTLs^48,49^ for variants located within a DHS and with a neighboring DHS in haplotype-resolved GM12878 Fiber-seq data. Across all 17 variants that fit these criteria, Mola predicted the correct direction of the co-accessibility effect at 14 loci, and predicted effect sizes were significantly correlated with the corresponding Fiber-seq measurements (**Figure S23**). For example, the caQTL rs59769368 provided a particularly clear experimental validation of this capability. Mola predicted strong coupling between the C reference allele at the variant-centered region and a distal regulatory region approximately 1.5–1.8 kb away, whereas this coupling was markedly reduced for the G alternative allele (**Figure 7G**). Haplotype-resolved GM12878 Fiber-seq molecules showed consistent changes in the co-accessibility measurement (**Figure 7G**). Mola therefore predicts not only the mean accessibility changes of the variant effect, but also the specific distal molecular relationship altered by the variant. Although the available experimental set was necessarily small, this agreement provides direct evidence that Mola can predict variant-driven changes in single-molecule regulatory coordination.

These analyses advance noncoding-variant prediction by shifting from mean regulatory activity to single-molecule nucleosome organization and coordination. By predicting allele-specific distributions, Mola identifies variants that couple or decouple regulatory elements, and generates testable hypotheses about long-range variant effects. This adds a new dimension to regulatory-variant prediction, directly linking sequence variation to single-molecule chromatin mechanisms.

## Discussion

We present a sequence-to-distribution (s2d) framework for directly learning single-molecule chromatin state distributions from DNA sequence. This framework and its Fiber-seq-trained model, Mola, define a new type of sequence model, shifting from predicting an average regulatory profile to predicting a distribution that captures heterogeneity across individual molecules. A major portion of chromatin heterogeneity and coordination can therefore be predicted from DNA sequence alone, indicating that this information is encoded in the sequence. Because every generated molecule is a sample from a sequence-conditioned distribution, Mola functions as “*virtual chromatin*”: a sequence edit becomes a controlled *in silico* experiment whose readout is a population of single molecules.

Large-scale *in silico* TF screening with Mola shows that TFs differ not only in whether they open chromatin but in how they organize the surrounding nucleosomes. Chromatin opening, nucleosome phasing, repeat length, and array periodicity can be separable, TF-dependent properties: CTCF showed strong opening, phasing, and local array remodeling, reducing nucleosome repeat length while largely preserving regularity. Cell-type specific factors like IRF and BATF decreased both spacing and periodicity, and ZNF146 emerged as a nucleosome-trapping factor that increases local nucleosome occupancy while phasing the array. These single-molecule signatures likely reflect the remodelers and cofactors each TF engages^31,32^, and we envision them as a scalable way to assign TFs to mechanistic classes in the future. The promoter analyses show how the model’s findings can be validated experimentally. Mola found YY1-associated promoters to have the strongest +1 nucleosome positioning, and experimental YY1 degradation data fully validate this conclusion^37^.

Mola also revealed that sequence encodes coordination between distal elements. The synthetic two-DHS screen showed that distal co-accessibility is TF-specific and not solely explained by local accessibility. Co-accessibility and co-binding largely shared the same motifs, and a plausible mechanism is TF hubs, local environments of high TF concentration that favor both TF binding and chromatin opening on the same molecule^41^. Joint modeling of accessibility and CpG methylation revealed sequence-based bimodality at a small subset of regulatory elements, aligning with a feedback model in which methylation-sensitive TF occupancy^43^, chromatin accessibility, and methylation maintenance stabilize alternative states, offering a candidate mechanism for sustaining epigenomic memory.

Beyond the observed distributions, Mola can also predict what the ground-truth data cannot answer directly. For example, how chromatin states change over time, and how genetic variants reshape them. Pseudo-dynamic trajectories revealed a clear hierarchy of time scales: TF footprints switched most rapidly, DHSs opened and closed much more slowly, and nucleosomes showed constrained, subdiffusive repositioning along DNA. Although frames are not calibrated to biological time, this hierarchy is consistent with rapid TF exchange in living cells^1^, slower accessibility loss after remodeler inhibition^44,45^, and confined nucleosome diffusion in vitro^46^. For variant effect prediction, Mola performed on par with or better than expert models on QTL effects on mean accessibility. Mola can also predict variant-dependent changes in co-accessibility and nucleosome array organization, which agreed with allele-resolved single-molecule data^16,49^.

Looking forward, we envision broad applications across areas. For chromatin biology, Mola serves as an engine for *in silico* single-molecule experiments, generating hypotheses and prioritizing perturbation experiments, as in the YY1 case. For single-molecule genomics, generated molecules provide a sequence-based expectation of co-accessibility at loci with few reads and can guide where deep sequencing is most informative. For human genetics, Mola provides a new score of variant effects on heterogeneity and coordination that current models do not, and that could be crucial for variants with little average effect. For sequence design, single-molecule heterogeneity can now become a design target, so that a sequence element can be designed to achieve a desired distribution.

Several limitations define the framework’s current scope. First, Mola was trained on Fiber-seq data from a single human lymphocyte context, so the learned distribution reflects the sequence, trans-environment, and cell state of the training data, and a sequence-only input does not mean chromatin state depends solely on sequence. Second, Fiber-seq offers a detailed but assay-specific view, limited by molecule depth, uneven nucleotide observability, underdetection of TF footprints, and lower m5C measurement quality. Third, the 10 kb window limits analysis of longer-range regulation.

These limitations also suggest clear directions for future studies. Training across cell types, developmental stages, and perturbations could distinguish general cis-regulatory rules from context-specific trans effects. Longer reads and larger datasets can reveal larger regulatory neighborhoods and resolve rare states. Additional single-molecule modalities, such as histone modifications and transcription on the same fiber, can be modeled jointly, and haplotype-resolved data across many individuals^16^ would allow single-molecule variant effects to be evaluated at scale.

In summary, the sequence-to-distribution framework and Mola advance regulatory genome modeling by moving beyond average chromatin signal prediction to capturing the probabilistic distribution of molecular states encoded by sequence. Beyond chromatin, this framework can more generally apply to biological properties better described as distributions, including variation among individual DNA and RNA molecules, among cells, and even among individuals. Sequence-to-distribution modeling provides a foundation for exploring the sequence basis of heterogeneity and dynamics of genome regulation, and how genetic variation changes it, one molecule at a time.

## Methods

### Model design

Mola (Model of single-molecule long-read chromatin accessibility) is a sequence-to-distribution model designed for modeling single-molecule regulatory genomics data like Fiber-seq. Given an input DNA sequence, it predicts the distribution of single-molecule chromatin states by generating any possible chromatin molecule conditioned on the underlying sequence. We formulated single-molecule chromatin state generation as a denoising diffusion-based generative model. The training target is a clean molecule *x*_0_, and the condition is the one-hot encoded DNA sequence *s* of the same window. A neural denoiser is trained to recover *x*_0_ from a noise-corrupted copy at any noise scale *σ*, given *s*. Because a denoiser trained in this way estimates the conditional score *∇_x_* log *p*_σ_(*x* | *s*), it implicitly defines a distribution over single-molecule states for a given sequence. New molecules are drawn from this distribution by integrating the reverse diffusion process from pure noise, and the neighborhood of a given chromatin state can be explored by running MCMC such as Langevin dynamics at a fixed noise level.

Two key properties of this model design enable the analyses and discoveries. First, the model generates long-read single-molecule chromatin states, enabling representation not only of mean accessibility and methylation but also of their covariation along a chromatin fiber, which underlies the co-accessibility, co-footprinting, and accessibility/methylation analyses. Second, the model is conditioned only on DNA sequence. Because sequence is upstream of chromatin state and is not altered by it, this conditioning modeling follows the causal direction of gene regulation, and editing the input sequence is the computational analog of a genetic perturbation. The model can therefore analyze and evaluate the impact of local sequence changes on chromatin states, including motif insertions, motif disruptions, and single-nucleotide variants.

### Training data

We trained Mola on a deep Fiber-seq dataset from GM12878 lymphoblastoid cells^16^ (BioProject PRJNA1233341). Reads were aligned to GRCh38 with pbmm2 v1.10.0 using the CCS preset. We used the aligned BAM containing m^6^A predictions from Fibertools-rs v0.1.4 and CpG methylation calls from Primrose v1.4.0. We excluded chromosomes 8, 9, and 10 from training and reserved them for testing and evaluation. Each generated molecule consists of two base-resolution tracks over the 10 kb window: exogenous m^6^A modification at A/T positions, which reports chromatin accessibility as measured by Fiber-seq, and endogenous CpG methylation at CpG sites, which reports DNA methylation. We refer to an observed or generated 10 kb single-molecule profile as a chromatin fiber.

Training examples were 10 kb windows. We sampled windows in proportion to a genome-wide weighting track derived from mean accessibility across molecules. Within each window, only molecules spanning the full 10 kb were retained. Fibers from accepted windows were concatenated into batches of variable size.

For each fiber, m^6^A and CpG modification likelihoods were placed at their aligned reference positions, with CpG calls strand-adjusted to the corresponding reference CpG. Missing values and positions without a detected modification signal were assigned a value of zero. This will provide us with a target likelihood tensor of dimensions N × 2 × 10,000 for N fibers. The DNA sequence was one-hot encoded in A, C, G, T order and repeated across the fibers from the same window. During training, the network therefore received six input channels: the four sequence channels and the two noise-corrupted signal channels.

### Model architecture and training

The denoising network was a one-dimensional U-Net. An initial convolutional block projected the six input channels to 512 features. All convolutional blocks used a kernel size of 17, group normalization, and GELU activation, with residual blocks in both the encoder and the decoder. Four encoder stages downsampled by factors of 4, 4, 5, and 5, yielding 25 bins of 400 bp at the bottleneck, where three additional 512-channel convolutional blocks were applied. The decoder mirrored the encoder, with linear upsampling factors of 5, 5, 4, and 4 and skip connections from the corresponding encoder stages. A final convolution projected the 512 features to the two output channels.

The network was conditioned on the noise level through t = 0.25 log(σ), where σ is the diffusion noise scale. A sinusoidal embedding of t was passed through a multilayer perceptron with SiLU activation, and learned projections of the resulting embedding were added to each downsampling and upsampling block.

We trained Mola with the preconditioning and weighted denoising objective of Karras et al.^51^ At each optimization step, an independent noise scale was drawn for each fiber:

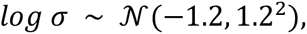

The clean target x_0_ was then corrupted with Gaussian noise:

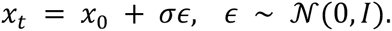

The preconditioning rescaled the noisy chromatin channels before they entered the network and combined the network output with a noise-dependent skip connection to form the estimate of the clean fiber:

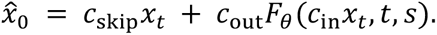

Here F_θ_ is the U-Net, t is the noise-level embedding input, and s is the DNA sequence. The scaling coefficients used a fixed reference data scale of σ_data_ = 0.3:

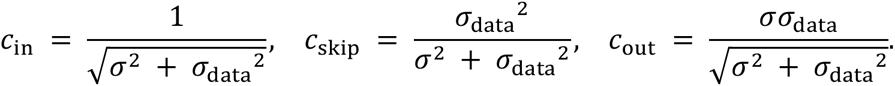

This scaling kept the network input at a comparable magnitude across noise levels. At low noise the estimate relied mainly on the noisy input itself, whereas at high noise it relied mainly on the learned prediction. The training loss was a weighted mean squared error (MSE) between the estimated and clean fibers:

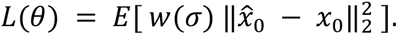

The weight 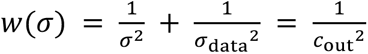 compensated for the output scaling. We averaged losses over channels and genomic positions, then over fibers in the batch. We trained with Adam at a learning rate of 1 × 10⁻³ on 4x NVIDIA H100 GPUs. As data augmentation, we reverse-complemented the input sequence with probability 0.5; in that case, we also reversed the target signals along the genomic axis. We trained for approximately 3 million steps, each on a batch of two 10 kb windows drawn in proportion to the mean-accessibility track, using reads that fully spanned each window (on average 100 molecules per batch), which amounts to roughly 300 million 10 kb read segments, or about 10 passes over the entire dataset.

### Model sampling (inference)

To generate fibers for a sequence s, we integrated the reverse diffusion process from Gaussian noise using the trained denoiser. The conditional score s_θ_(x, σ | s) = ∇_x_ log p_σ_(x | s) was estimated from the denoiser as:

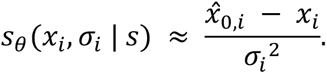

Here x_i_ is the noisy fiber and x^_0,i_ is the predicted clean fiber at noise scale σ_i_. All fibers generated for a region shared the same sequence encoding, but each was initialized from independent Gaussian noise with variance σ_max_^2^. We used the deterministic sampler of a 10-step Karras schedule with σ_min_ = 0.002, σ_max_ = 80, and ρ = 7. The nonzero noise levels at normalized schedule position r_i_ were:

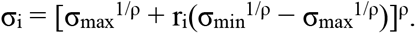

A final zero-noise step completed the schedule. At each step, the noisy fiber was updated by Euler integration:

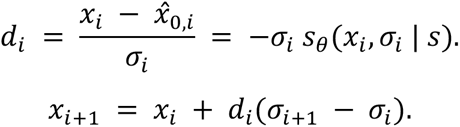

All fibers generated for a region were conditioned on the same sequence encoding and differed only in their initial noise. Given that noise, the sampler is deterministic, so each sample is one fiber from the learned conditional distribution. A generated fiber consists of two continuous tracks on the scale of the training targets, m6A at each base and CpG methylation at each CpG; values at bases where the modification cannot occur (m6A at C or G, methylation outside CpGs) were ignored. Unless stated otherwise, we generated all fibers in this study with this 10-step Euler sampler.

### Model evaluation

We evaluated Mola on the held-out chromosomes 8, 9, and 10 using 10 kb windows centered on GM12878 DNase I hypersensitive sites (DHSs; ENCODE ENCFF762CRQ). Regions were required to have at least 20 observed molecules spanning the full window. For each region, we generated 500 fibers, regardless of the observed coverage. Accessibility was evaluated only at reference A/T positions, and DNA methylation only at reference CpGs.

We calculated mean accessibility profiles across fibers at each A/T position. We defined co-accessibility as the Pearson correlation between binned signals across fibers; representative co-accessibility maps used 20 bp bins. We measured agreement between generated and observed maps as the mean correlation between corresponding rows of the matrix. Mean CpG methylation profiles were calculated across fibers at each valid CpG and compared between generated and observed fibers by Pearson correlation.

Because both generated and observed statistics are estimated from a finite number of fibers, we estimated the split-half reliability of each set separately. Each set was divided into two equally sized, consecutive subsets, the relevant statistic was recomputed in each half, and the correlation between halves was corrected with the Spearman–Brown formula:

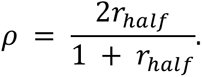

The resulting reliabilities ρ_model_ and ρ_data_ define a statistical correlation upper bound (ceiling) r_max_ = √(ρ_model_ ρ_data_), an estimate of the maximum generated-versus-observed correlation attainable given the finite-sample noise in both sets. The efficiency was the generated-versus-observed correlation divided by r_max_.

### *In silico* motif screening

The motif library comprised 285 nonredundant transcription factor (TF) motifs selected from the archetype seed motifs of Vierstra et al.^20^, with one representative per consensus sequence. For each motif and genomic background, we replaced the corresponding reference bases with the forward consensus sequence within a 10 kb background window, then generated fibers for the edited and unedited sequences with 10 Euler steps. Each condition was sampled with 100 fibers in each of 100 DHS or random backgrounds; the single-copy screen was also performed using 100 DHS backgrounds from chromosome 10. Multicopy insertions used 50 bp start-to-start spacing (e.g., first motif starts at 0 bp, second motif starts at 50 bp, third motif starts at 100 bp). Effects were calculated against the unedited sequence within each background before averaging across backgrounds.

### Nucleosome opening and phasing

Nucleosomes on each fiber were called with FiberHMM^40^. For this purpose, we represented generated fibers as 10 kb pseudo-reads carrying the edited reference sequence. At canonical A/T positions, m^6^A predictions greater than 0.5 were encoded as modified bases and the remaining A/T positions as unmodified. We ran FiberHMM in Hia5/PacBio mode with a modification-probability threshold of 128 and a minimum read length of 1,000 bp. Nucleosome positions were extracted from the resulting BAM and converted to per-base occupancy profiles.

For each motif and background, nucleosome occupancy was averaged across fibers. We defined opening power as one minus the mean occupancy in the motif interval extended by 50 bp on each side; in the single-copy screen, this interval was from 4,950 to 5,050 + L, where L is the motif length. The opening effect was the opening power of the edited sequence minus that of the unedited control, so that positive values indicate reduced nucleosome occupancy relative to the background.

Nucleosome phasing was quantified from the strength of regularly spaced peaks in mean nucleosome occupancy flanking each inserted motif. Within each background, we averaged occupancy across fibers, detrended the 2 kb regions on either side of the central opening interval, and multiplied them by a Hann window to minimize boundary artifacts. We computed the Fourier spectrum of each flank, averaged the spectra across backgrounds, and then selected the strongest component with a period of 160–210 bp on each side. We defined the phasing score as the mean amplitude across the two flanks, and the phasing effect as the score after motif insertion minus the unedited background score; positive values indicate a stronger, more regularly phased nucleosome pattern.

For ZNF146 validation, we aggregated observed GM12878 Fiber-seq profiles around chromosome-10 ZNF146 ChIP-seq peaks from K562 cells (ENCODE ENCFF932FVX) and compared them with the *in silico* insertion result.

### Nucleosome array analyses

Nucleosome array spacing and regularity were measured from the accessibility signal in the 1 kb region at positions 5,200–6,200 bp, adjacent to the motif insertion center at 5,000 bp. For each molecule, we computed the autocorrelation of the accessibility profile, that is, its correlation with copies of itself shifted by increasing lags; when the lag matches the spacing of a repeating pattern, the shifted profiles align and produce a peak. We normalized each autocorrelation curve to 1 at zero lag and averaged first across molecules within a background, then across backgrounds. After smoothing the averaged curve, we identified the highest peak and refined its position and height with a quadratic fit through the peak and its two neighboring points. We defined the nucleosome repeat length (NRL) as the peak position in base pairs, an estimate of the typical distance between neighboring nucleosomes. We defined nucleosome periodicity strength (NPS) as the height of the same peak above zero, with higher values indicating a more regular repeating pattern.

### Promoter +1 nucleosome analyses

Promoter annotations and motif-contribution scores were obtained from the Puffin study^35^. Puffin defines the contribution of each motif as its position-dependent effect averaged within ±20 bp of the transcription start site (TSS), weighted by predicted transcription initiation. We used the top 16,000 human TSSs ranked by initiation signal, assigned each a dominant motif by its contribution score, and thereby grouped promoters into eight motif classes: TATA, YY1, SP, ETS, NFY, CREB, NRF1, and ZNF143. On chromosomes 8–10, 1,585 promoters belonged to these eight classes, including 89 TATA-dominant and 88 YY1-dominant promoters. We sampled each promoter and condition from 100 fibers using 10 Euler steps. Minus-strand profiles were reversed after binning so that all profiles are displayed in the direction of transcription.

To test the effect of YY1 in TATA promoters, the 12 bp YY1 motif sequence CAAAATGGCGGC replaced positions 5,020–5,032 of the reference-oriented 10 kb window in each of the 757 TATA-dominant promoters genome-wide (all chromosomes, including those used for training). We used the same genomic orientation and offset for both strands, so the insertion lay downstream of plus-strand TSSs and upstream of minus-strand TSSs. Conversely, we depleted YY1 *in silico* from each of the 777 YY1-dominant promoters genome-wide by randomizing the 20 bp intervals centered on its highest-scoring YY1 site, replacing each base with one of the other three nucleotides with equal probability. We sampled original and mutated promoters in matched pairs.

Within the central 2 kb (positions 4,000–6,000), the m^6^A signal was masked to reference A/T positions, averaged into 10 bp bins, and then averaged across fibers to give each promoter’s accessibility profile. For display, group means and standard errors were smoothed with a two-bin Gaussian kernel and scaled by the maximum smoothed mean within each condition. Co-accessibility was calculated across fibers for every pair of bins in the central 1 kb (−500 to +500 bp from the TSS) and averaged with equal weight across promoters. We quantified the +1 nucleosome strength of each promoter by the depth of the accessibility minimum at the +1 nucleosome position relative to the flanking signal. Group values are means with 95% bootstrap confidence intervals over promoters. Differences among the eight promoter groups were tested with the Kruskal–Wallis test; paired comparisons (YY1 insertion versus original TATA promoters, YY1 mutation versus wild-type promoters, and each degron time point versus 0 h) used two-sided Wilcoxon signed-rank tests on per-promoter differences.

As an experimental comparison for the YY1 depletion, we used ATAC-seq from KOLF2.2J human induced pluripotent stem cells at 0, 6, and 24 h after auxin-induced YY1 degradation (GEO GSE343368), with two replicates per time point. For this comparison, TATA- and YY1-dominant promoters were selected from the same TSS table across all chromosomes.

### Co-accessibility between DHSs

To screen for co-accessibility between regulatory element pairs, we constructed synthetic DHS pairs in genomic backgrounds. Each synthetic element contained three homotypic copies of a TF motif at offsets of 0, 50, and 100 bp. The anchor element began at position 1,000, and the second element was placed 1–8 kb downstream. For each TF pair, distance, and background, we generated 200 fibers. Single-molecule accessibility profiles were computed at A/T positions of the edited sequence and averaged into 20 bp bins, and correlations were calculated across the 200 fibers for every pair of anchor and distal bins within each background and TF pair. For validation, we identified motif-matched pairs of GM12878 DHSs within 10 kb of each other and used observed molecules spanning both elements. Correlations were calculated as above, and the observed and generated TF-by-TF matrices were compared by Pearson correlation. To test whether a partner DHS changes the accessibility of the target DHS, we compared the mean A/T accessibility in the target window (−60 to +200 bp from the target insertion start, excluding the inserted motif spans) between the two-site condition and the target-only control, pooling distances of 1–8 kb and both placements of partner and target. We shrunk the effect toward zero by one standard error estimated from the eight distance-specific effects, and the resulting partner-by-target matrix is shown in the shared TF-family order of Figure 4.

### TF co-footprinting

To test whether two TFs occupy the same molecule, we converted generated fibers to pseudo-reads and used FiberHMM to call TF footprints, nucleosome-sized protected segments, and methyltransferase-sensitive patches (MSPs, open regions). A motif was called bound if a single TF block of at most 60 bp covered at least 75% of its A/T positions, and unbound if no TF block covered 25% or more; intermediate cases were called ambiguous. We further required both motif copies (anchor and distal) to lie within MSPs on the same pseudo-read and excluded ambiguous calls and broad footprints that could not be assigned to a single copy.

For each background, distance, and TF pair, molecules were classified as bound, anchor only, distal only, or neither bound. We calculated the binary co-binding correlation by summing the within-background covariance and variance components across strata before normalization. Confidence intervals were estimated from 2,000 bootstrap resamples of genomic backgrounds, and the final co-binding correlations were shrunk toward zero by one standard error: sign(r) × max(|r| − SE_bootstrap_, 0).

### Accessibility and CpG methylation

Because each fiber carries both an accessibility track and a CpG methylation track, we examined how the two vary together on individual molecules. For each molecule, we computed the mean A/T accessibility over all reference A/T positions in the DHS and the methylated-CpG fraction, defined as the fraction of reference CpGs with a methylation signal of at least 0.5. Within each DHS, we calculated the Pearson correlation between these two per-molecule values across molecules (the cross-molecule correlation); across DHSs, we correlated the DHS-level means (the across-DHS correlation). The genome-wide across-DHS analysis used 66,033 DHS-centered 1 kb windows containing at least five CpGs and at least 50 molecules.

To identify DHSs with two methylation states, we modeled per-molecule methylated-CpG counts with beta-binomial mixtures. For each observed molecule, k was the number of called reference CpGs with a methylation probability of at least 0.5, and n was the total number of called reference CpGs. Molecules were retained when they had at least five called CpGs covering at least 80% of the reference CpGs in the DHS, and a DHS was tested when it had at least five reference CpGs and at least 50 passing molecules. We fitted one- and two-component beta-binomial mixtures, constrained α and β of each component to be at least 1, used four random initializations, and compared the two fits with a sequential parametric bootstrap likelihood-ratio test. A DHS was called two-state (bimodal) when q < 0.05, the fitted component means differed by at least 0.25, and the minor component contained at least 10% of molecules. To ask whether sequence predicts which DHSs are two-state, we computed two scores from generated molecules only: the standard deviation across molecules of the methylated-CpG fraction, and the negative of the cross-molecule accessibility–methylation correlation. Each score ranked the held-out DHSs, and performance was summarized by the area under the ROC curve for separating observed two-state from unimodal DHSs (28 versus 2,702 DHSs with complete scores), with 95% confidence intervals from 2,000 class-stratified bootstrap resamples.

For motif enrichment, each genome-wide two-state DHS was matched one-to-one within its chromosome to a unimodal DHS. For each motif, we scored the centered 1 kb sequence of each DHS by its maximum log2-odds match and compared the paired differences with one-sided Wilcoxon signed-rank tests, Benjamini–Hochberg corrected across archetypes.

In the motif-insertion screen for accessibility–methylation coupling, four copies of each of the 285 motifs were inserted 50 bp apart into 20 backgrounds with 100 generated molecules per background, and the Pearson correlation between per-molecule mean A/T accessibility and mean CpG methylation was computed over all molecules. We tested NRF1 dosage with one to six copies spaced 50 bp apart in the same 20 backgrounds with 100 molecules each, and evaluated the joint distribution of accessibility and methylation within 600 bp of the insertion site.

### Simulating pseudo-dynamic trajectories

We used sequence-conditioned Langevin sampling to explore transitions between chromatin configurations supported by Mola. We generated 100 independently seeded trajectories for the 10-kb NAPA region (GRCh38 chr19:47,509,000–47,519,000; zero-based, half-open coordinates). Initial fibers were generated with the 10-step Euler sampler and perturbed with Gaussian noise of standard deviation 0.9.

The chain was run at a fixed noise level of σ = 0.9. For noisy state xₖ and DNA sequence s, the denoiser Dθ estimates the clean state and provides an estimate of the conditional score:

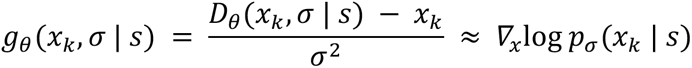

Here pσ is the sequence-conditioned distribution after Gaussian noise is added at scale σ. At each update, we rescaled the estimated score when its root-mean-square magnitude exceeded 10, giving the clipped score s̃ₖ. The Langevin update combined this score-directed drift with independent Gaussian noise:

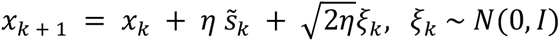

The drift favors locally higher-density configurations, while the stochastic term permits exploration of alternative states. We used step size η = 0.01 and a Gaussian-noise multiplier of 1.0, without momentum or a Metropolis acceptance step. The fixed σ sets the noise level at which we evaluate the score; the stochastic increment in each update is scaled by the square root of 2η.

After 5,000 initial updates, we saved one frame after each of the next 10,000 updates. Each saved noisy state was evaluated once with the denoiser at σ = 0.9 to obtain the clean-state readout. This readout did not replace the noisy state used for subsequent Langevin updates. We applied readout clipping and rounding downstream for footprint calling and visualization. For FiberHMM calling, we used the Hia5/PacBio model, a threshold of 128, and a minimum read length of 1,000 bp, retaining 85–250-bp nucleosome footprints.

We classified CTCF-associated footprint states at chr19:47,515,580–47,515,594 using motif protection and surrounding DNA accessibility. A candidate footprint was a FiberHMM-called protected block ≤60 bp long that covered at least three of the motif’s four A/T positions and lay within a methyltransferase-sensitive (accessible) patch. Frames were analyzed only when the pooled mean A/T accessibility in the two flanking windows, 5–40 bp outside the motif boundaries, was ≥0.5 and no single eligible nucleosome footprint covered ≥75% of the motif. Within these frames, ON required a candidate footprint and mean motif A/T accessibility ≤0.25; OFF required no candidate footprint and mean motif A/T accessibility ≥0.5. Frames with insufficient flank accessibility or excessive nucleosome overlap were classified as unresolved, rather than OFF.

For the 193-bp DHS (chr19:47,518,230–47,518,423), we measured the fraction of DHS bases covered by one or more eligible nucleosome footprints, counting each base only once where footprints overlapped. The DHS was classified as closed when coverage was ≥75%, and open when coverage was ≤25% and mean accessibility across DHS A/T bases was ≥0.5.

For both CTCF footprint and DHS states, confirmation required five consecutive frames meeting the relevant criteria; state onset was assigned to the first of those five frames. We estimated 95% confidence intervals from 2,000 bootstrap resamples of whole-molecule trajectories.

For nucleosome diffusion estimates, nucleosome footprints were greedily linked between adjacent frames using a 20-bp center-displacement limit and a matching cost of absolute center displacement plus 0.1 times absolute length change, without frame gaps. We computed mean absolute displacement and mean-squared displacement (MSD) from continuously linked endpoint pairs, averaged within molecules and then equally across eligible molecules. Root-mean-square displacement was the square root of MSD. Displacement confidence intervals used 5,000 whole-molecule bootstrap resamples.

We fitted the following model, with nonnegative offset C, to MSD at lags of 1, 2, 5, 10, 20, 50, 100 and 200 frames:

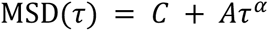

### Variant effects benchmark with dsQTLs and caQTLs

To predict variant effects, we generated fibers for the reference- and alternative-allele 10 kb sequences, which differed only at the variant position. We sampled both alleles with the same initial states, random noise, and schedule, so differences reflect the sequence change alone. For the genome-wide dsQTL and caQTL analyses, we sampled 100 fibers per allele and variant; the high-depth chromosome-17 example used 500 fibers per allele.

The mean-accessibility effect was log_2_[(μ_ALT_ + 10⁻⁶)/(μ_REF_ + 10⁻⁶)], where μ was the mean m^6^A prediction over all fibers within the 200 bp window centered on the variant. We compared predicted effects with measured effects using Pearson correlation for positive dsQTLs in Yoruba lymphoblastoid cells and positive caQTLs in African lymphoblastoid cells. The dsQTL classification benchmark used published labels for 27,373 Yoruba variants (560 positives and 26,813 negatives). We ranked variants by the absolute Mola-predicted log fold change and measured performance by average precision. For comparison, we evaluated precomputed log fold changes from the ChromBPNet ATAC and ChromBPNet DNase models on the same variants.

### Variant effects at the single-molecule level

Beyond mean accessibility, we examined predicted variant effects on single-molecule co-accessibility. The melanoma variant example compared the two-base CC and TT alleles at GRCh38 positions 19,447,245–19,447,246. For validation with haplotype-resolved GM12878 Fiber-seq data, candidate positive dsQTLs or caQTLs had to overlap an ENCFF762CRQ DHS and have a distinct neighboring DHS whose nearest edge was 1–5 kb from the variant, with at least three SNP-covering reads per allele. The local region spanned −100 to +100 bp from the variant, and the distal interval covered the complete neighboring DHS. For each allele, co-accessibility was the mean correlation over all local–distal bin pairs, and the allelic effect was the allele-1 score minus the allele-2 score.

### Statistical analysis

Unless stated otherwise, correlations were Pearson coefficients. We measured classification performance by average precision or the area under the receiver operating characteristic curve, as specified for each benchmark. Bootstrap confidence intervals were percentile intervals unless stated otherwise.

## Data availability

GM12878 Fiber-seq data are available under BioProject PRJNA1233341, and processed FIRE data at Zenodo (doi:10.5281/zenodo.14511246). GM12878 DNase I hypersensitive sites (ENCFF762CRQ), ZNF146 ChIP-seq peaks from K562 cells (ENCFF932FVX), the HCT116 PRO-seq track (ENCFF621BFR), and the GM12878 H3K27ac (ENCFF469WVA) and H3K4me3 (ENCFF927KAJ) tracks were obtained from ENCODE. Puffin promoter annotations, motif-contribution scores, and aggregated PRO-cap tracks are available at https://github.com/jzhoulab/puffin_manuscript. Motif archetypes were obtained from the Vierstra motif-clustering resource (https://www.vierstra.org/resources/motif_clustering), which includes HOCOMOCO v11, JASPAR 2018, and HT-SELEX motifs. YY1-degron ATAC-seq data were obtained from GEO (GSE343368; GSM9949908–GSM9949913). QTL benchmarking tables and precomputed ChromBPNet variant scores were obtained from the ChromBPNet figure resources (https://github.com/kundajelab/chrombpnet-figures; Synapse syn64126763). The trained Mola checkpoint is available at Zenodo (doi:10.5281/zenodo.23108712).

## Code availability

Mola was implemented in Python with PyTorch. The code for model training, sampling, in silico perturbation, and all analyses in this study is available at https://github.com/jzhoulab/mola.

## Acknowledgements

We thank members of the Zhou laboratory for discussions and support. R.W. is the Illini 4000 Quantitative Biology Fellow of the Damon Runyon Cancer Research Foundation (DRQ-24-25). J.Z. is supported by the National Institutes of Health (NIGMS R35GM164319 and DP2GM146336, and NHGRI R01HG013724). S.P. is supported by the National Institutes of Health (NHLBI R01HL188168). This work used computational resources provided by the Center for Research Informatics (CRI) at the University of Chicago.

## Author contributions

R.W. and J.Z. conceived and designed the study, developed the computational methods, performed the analyses, and wrote the manuscript. J.J. and S.P. contributed to the analyses and to the writing. All authors read and approved the manuscript.

