## Supplementary Figures for "Predicting single-molecule chromatin distribution from DNA sequence"

### Figure S1

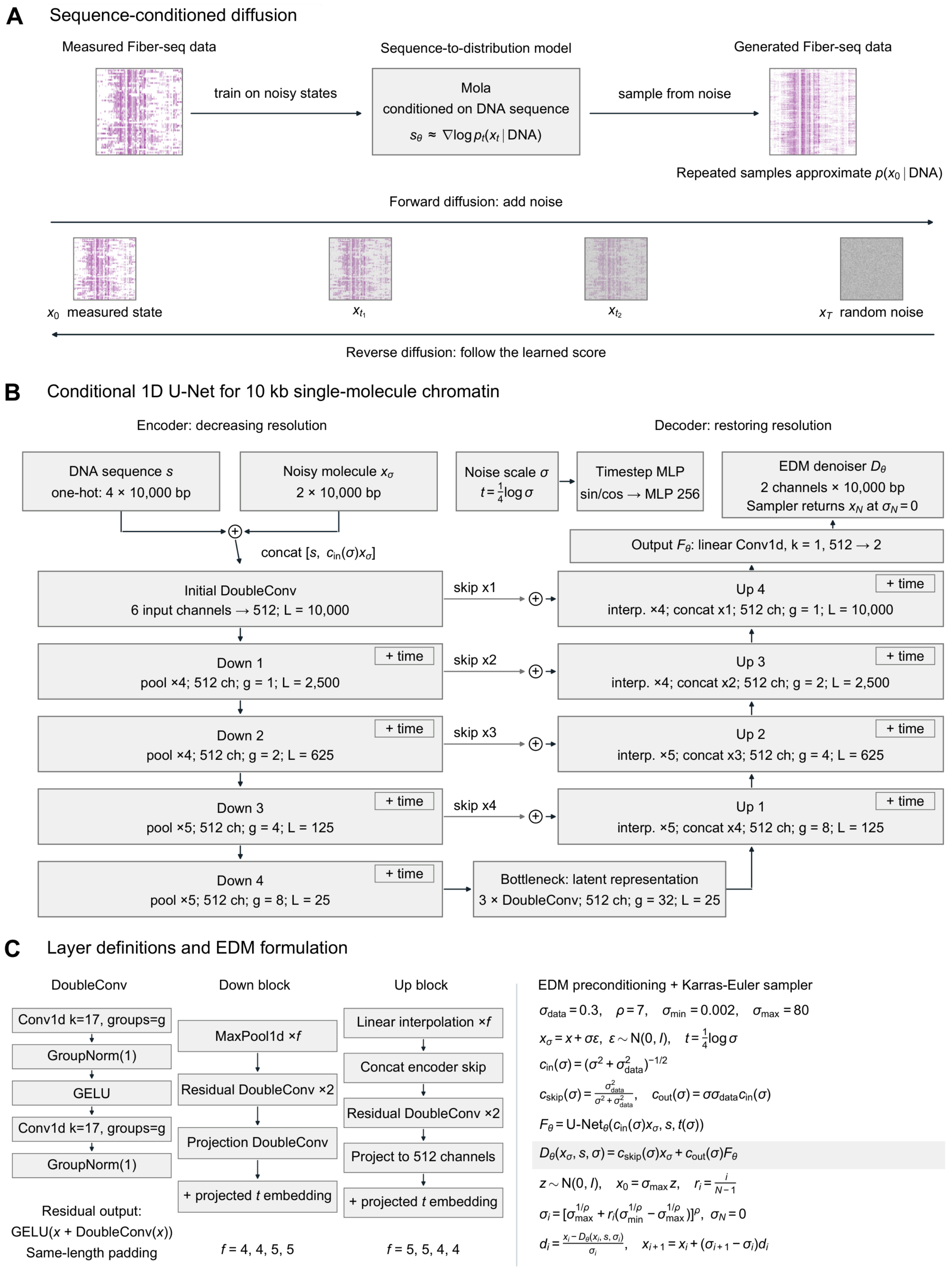

A
Sequence-conditioned diffusion
Measured Fiber-seq data
train on noisy states
Sequence-to-distribution model
Mola
conditioned on DNA sequence
$s_\theta \approx \nabla \log p_t(x_t \mid \text{DNA})$
sample from noise
Generated Fiber-seq data
Repeated samples approximate $p(x_0 \mid \text{DNA})$
Forward diffusion: add noise
$x_0$ measured state
$x_{t_1}$
$x_{t_2}$
$x_T$ random noise
Reverse diffusion: follow the learned score
B
Conditional 1D U-Net for 10 kb single-molecule chromatin
Encoder: decreasing resolution
Decoder: restoring resolution
DNA sequence $s$
one-hot: 4 × 10,000 bp
Noisy molecule $x_\sigma$
2 × 10,000 bp
Noise scale $\sigma$
$t = \frac{1}{4}\log\sigma$
Timestep MLP
sin/cos → MLP 256
EDM denoiser $D_\theta$
2 channels × 10,000 bp
Sampler returns $x_N$ at $\sigma_N = 0$
concat [$s$, $c_{in}(\sigma)x_\sigma$]
Output $F_\theta$: linear Conv1d, k = 1, 512 → 2
Initial DoubleConv
6 input channels → 512; L = 10,000
skip x1
Up 4
interp. ×4; concat x1; 512 ch; g = 1; L = 10,000
+ time
Down 1
pool ×4; 512 ch; g = 1; L = 2,500
+ time
skip x2
Up 3
interp. ×4; concat x2; 512 ch; g = 2; L = 2,500
+ time
Down 2
pool ×4; 512 ch; g = 2; L = 625
+ time
skip x3
Up 2
interp. ×5; concat x3; 512 ch; g = 4; L = 625
+ time
Down 3
pool ×5; 512 ch; g = 4; L = 125
+ time
skip x4
Up 1
interp. ×5; concat x4; 512 ch; g = 8; L = 125
+ time
Down 4
pool ×5; 512 ch; g = 8; L = 25
+ time
Bottleneck: latent representation
3 × DoubleConv; 512 ch; g = 32; L = 25
C
Layer definitions and EDM formulation
DoubleConv
Conv1d k=17, groups=g
GroupNorm(1)
GELU
Conv1d k=17, groups=g
GroupNorm(1)
Residual output:
GELU($x$ + DoubleConv($x$))
Same-length padding
Down block
MaxPool1d ×$f$
Residual DoubleConv ×2
Projection DoubleConv
+ projected $t$ embedding
$f$ = 4, 4, 5, 5
Up block
Linear interpolation ×$f$
Concat encoder skip
Residual DoubleConv ×2
Project to 512 channels
+ projected $t$ embedding
$f$ = 5, 5, 4, 4
EDM preconditioning + Karras-Euler sampler
$\sigma_{data} = 0.3, \quad \rho = 7, \quad \sigma_{min} = 0.002, \quad \sigma_{max} = 80$
$x_\sigma = x + \sigma\varepsilon, \ \varepsilon \sim N(0, I), \quad t = \frac{1}{4}\log\sigma$
$c_{in}(\sigma) = (\sigma^2 + \sigma_{data}^2)^{-1/2}$
$c_{skip}(\sigma) = \frac{\sigma_{data}^2}{\sigma^2 + \sigma_{data}^2}, \quad c_{out}(\sigma) = \sigma\sigma_{data}c_{in}(\sigma)$
$F_\theta = \text{U-Net}_\theta(c_{in}(\sigma)x_\sigma, s, t(\sigma))$
$D_\theta(x_\sigma, s, \sigma) = c_{skip}(\sigma)x_\sigma + c_{out}(\sigma)F_\theta$
$z \sim N(0, I), \quad x_0 = \sigma_{max} z, \quad r_i = \frac{i}{N-1}$
$\sigma_i = [\sigma_{max}^{1/\rho} + r_i(\sigma_{min}^{1/\rho} - \sigma_{max}^{1/\rho})]^\rho, \ \ \sigma_N = 0$
$d_i = \frac{x_i - D_\theta(x_i, s, \sigma_i)}{\sigma_i}, \quad x_{i+1} = x_i + (\sigma_{i+1} - \sigma_i)d_i$

**Figure S1. Architecture and training of the sequence-to-distribution model.** (A) Training corrupts observed GM12878 Fiber-seq molecules with Gaussian noise (forward diffusion); generation starts from noise and follows the learned, sequence-conditioned score (reverse diffusion), and repeated sampling approximates the distribution of chromatin states. (B) One-dimensional U-Net over a 10 kb window, conditioned on the one-hot sequence and the noise level ("+ time"). (C) Layer composition and EDM formulation with Karras–Euler sampling.

#### Figure S2

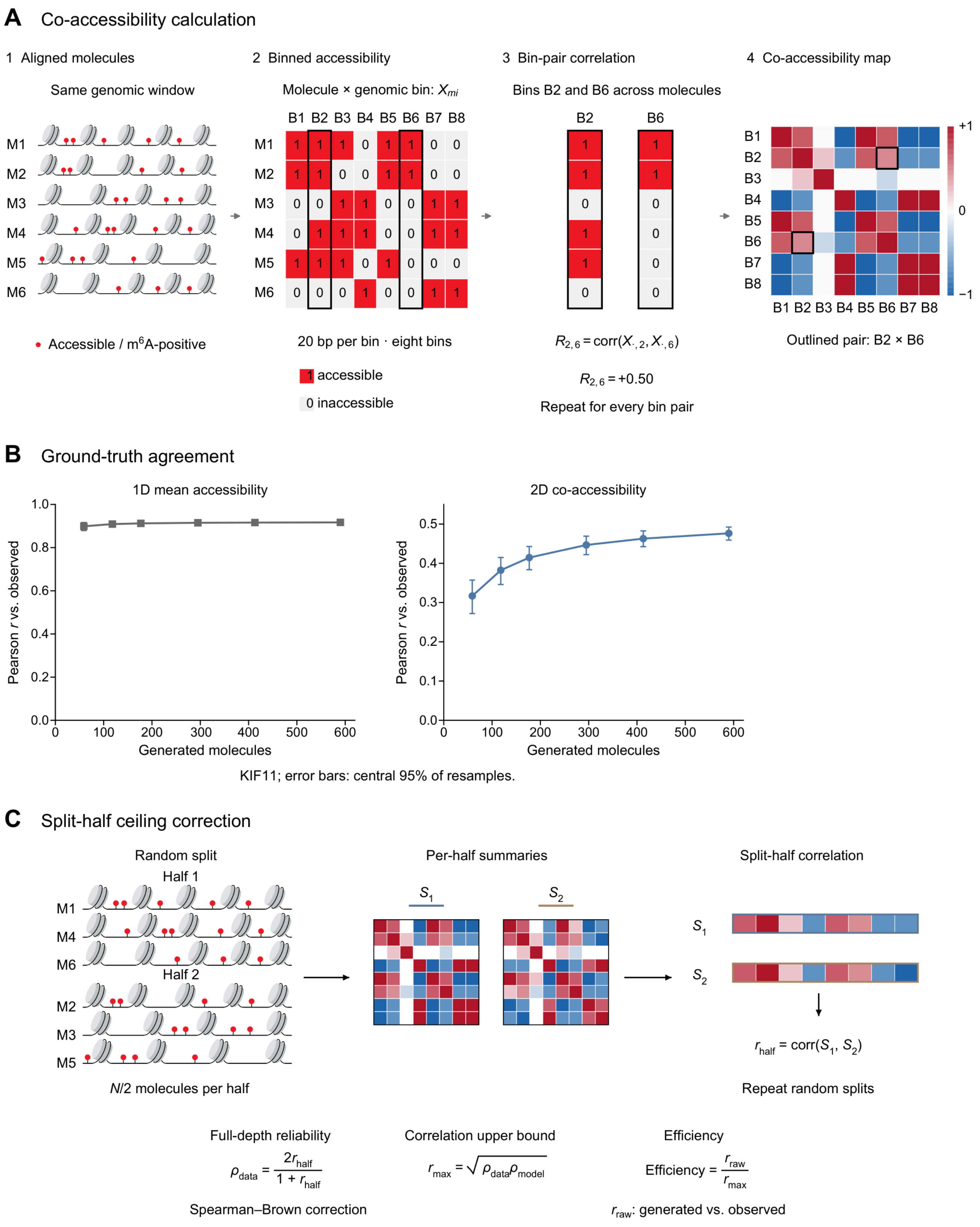

A Co-accessibility calculation
1 Aligned molecules
Same genomic window
M1
M2
M3
M4
M5
M6
Accessible / m6A-positive
2 Binned accessibility
Molecule × genomic bin: $X_{mi}$
B1 B2 B3 B4 B5 B6 B7 B8
20 bp per bin · eight bins
1 accessible
0 inaccessible
3 Bin-pair correlation
Bins B2 and B6 across molecules
B2
B6
$R_{2,6} = \mathrm{corr}(X_{\cdot,2}, X_{\cdot,6})$
$R_{2,6} = +0.50$
Repeat for every bin pair
4 Co-accessibility map
B1 B2 B3 B4 B5 B6 B7 B8
+1
0
−1
Outlined pair: B2 × B6
B Ground-truth agreement
1D mean accessibility
2D co-accessibility
Pearson r vs. observed
Generated molecules
KIF11; error bars: central 95% of resamples.
C Split-half ceiling correction
Random split
Half 1
Half 2
N/2 molecules per half
Per-half summaries
$S_1$
$S_2$
Split-half correlation
$r_{half} = \mathrm{corr}(S_1, S_2)$
Repeat random splits
Full-depth reliability
$\rho_{data} = \frac{2r_{half}}{1 + r_{half}}$
Spearman–Brown correction
Correlation upper bound
$r_{max} = \sqrt{\rho_{data}\rho_{model}}$
Efficiency
$\text{Efficiency} = \frac{r_{raw}}{r_{max}}$
$r_{raw}$: generated vs. observed

**Figure S2. Co-accessibility, sampling depth, and the split-half ceiling.** (A) Co-accessibility of two bins (20 bp) is the Pearson correlation of their accessibility across molecules. (B) Agreement with 59 observed molecules at the KIF11 promoter versus the number of generated molecules, for mean accessibility (left) and co-accessibility (right); points, means of 2,000 resamples; error bars, central 95%. (C) The split-half ceiling bounds model–data agreement; efficiency is the raw correlation divided by this ceiling. (A, C) Schematics.

### Figure S3

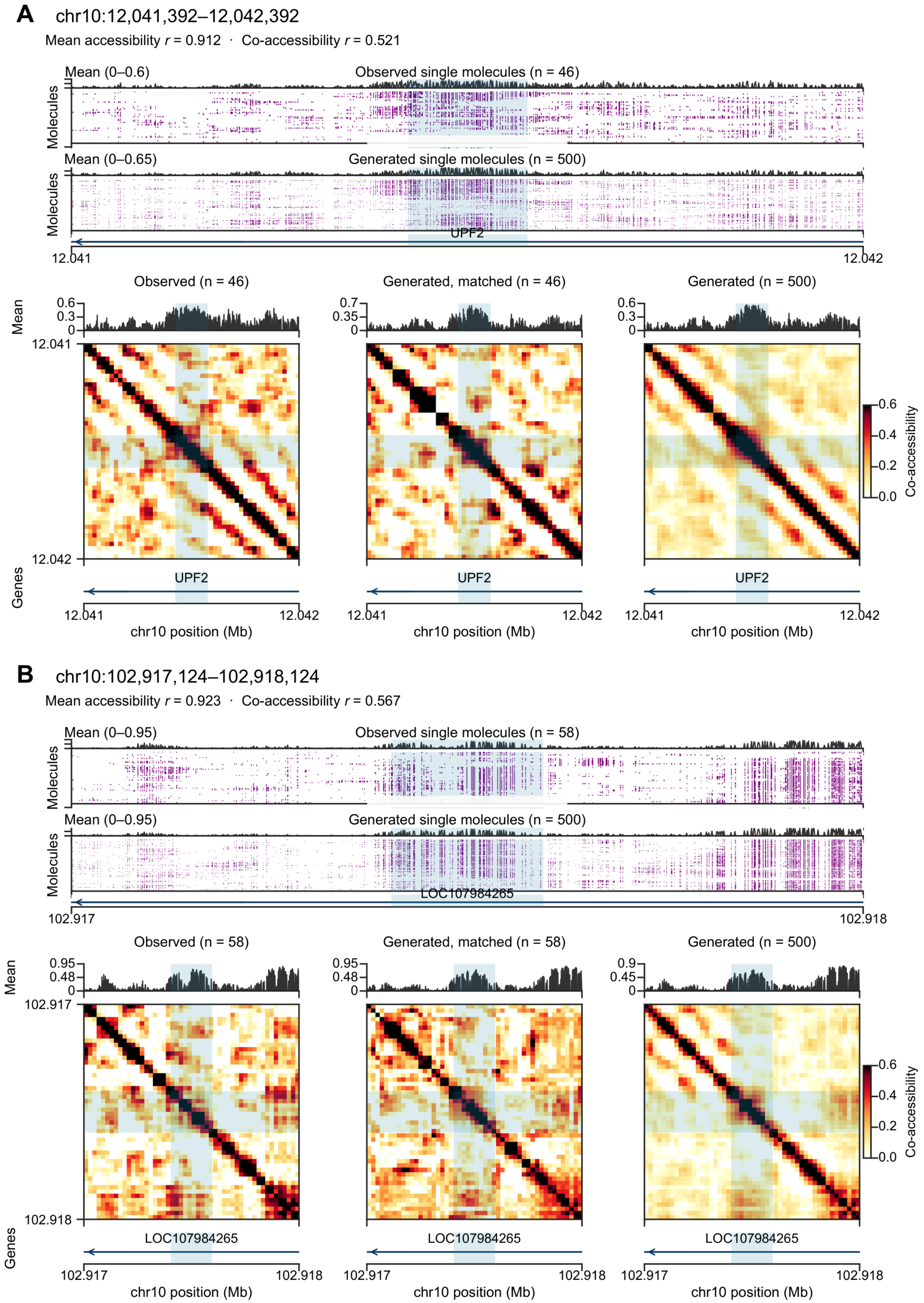

A
chr10:12,041,392–12,042,392
Mean accessibility r = 0.912 · Co-accessibility r = 0.521
Mean (0–0.6)
Observed single molecules (n = 46)
Molecules
Mean (0–0.65)
Generated single molecules (n = 500)
UPF2
12.041
12.042
Observed (n = 46)
Generated, matched (n = 46)
Generated (n = 500)
Mean
0.6
0.3
0
0.7
0.35
Co-accessibility
0.0
0.2
0.4
0.6
Genes
chr10 position (Mb)
B
chr10:102,917,124–102,918,124
Mean accessibility r = 0.923 · Co-accessibility r = 0.567
Mean (0–0.95)
Observed single molecules (n = 58)
Generated single molecules (n = 500)
LOC107984265
102.917
102.918
Observed (n = 58)
Generated, matched (n = 58)
0.95
0.48
0.9
0.45

**Figure S3. Held-out chromosome 10 examples of single-molecule accessibility and co-accessibility.** Two 1 kb windows centered on DHSs (shaded): observed and generated molecules with mean tracks, and co-accessibility maps (20 bp bins) from observed, depth-matched generated and all 500 generated molecules. Agreement with observed data is given above each example.

### Figure S4

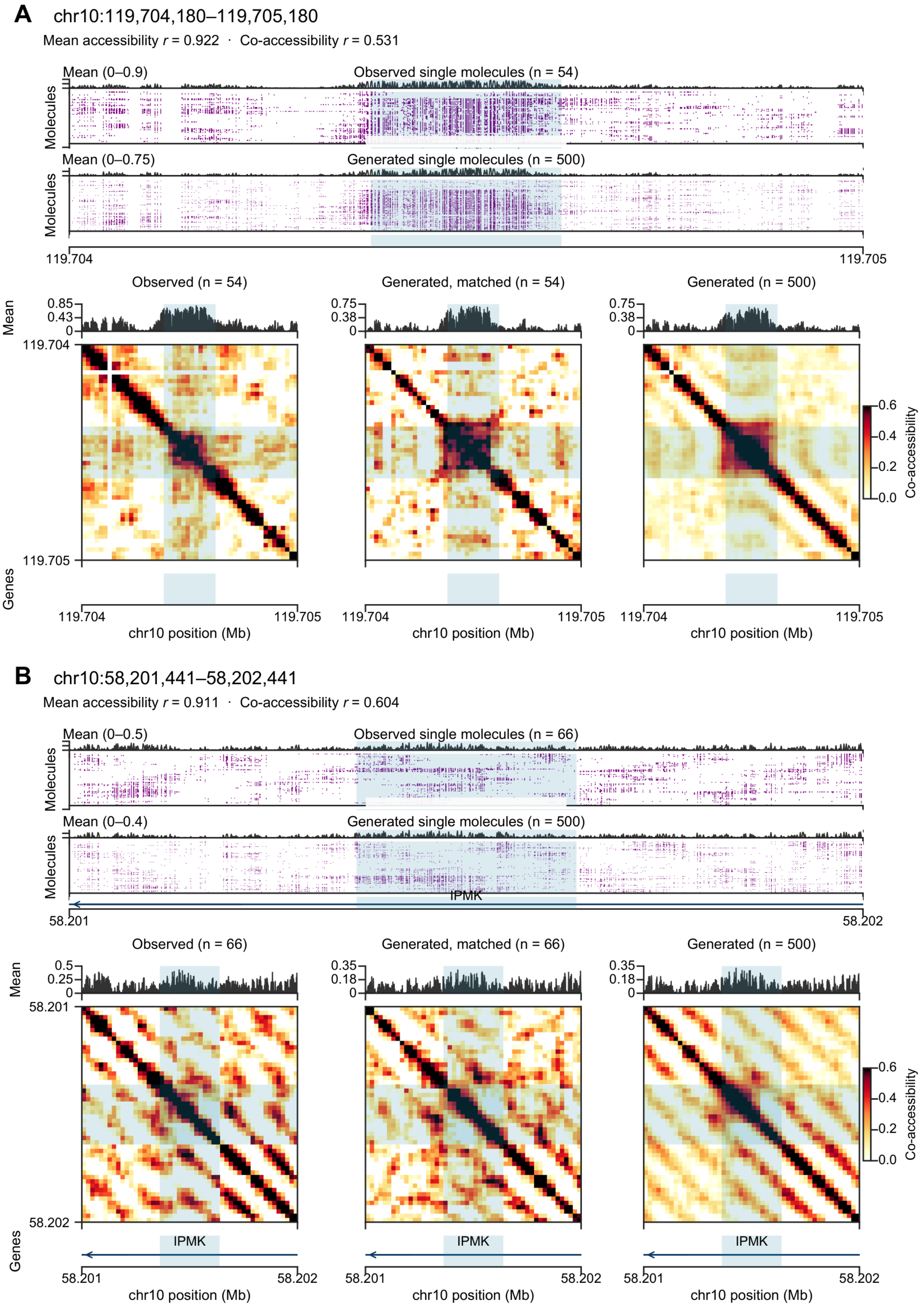


**Figure S4. Further held-out chromosome 10 examples.** As in Figure S3.

### Figure S5

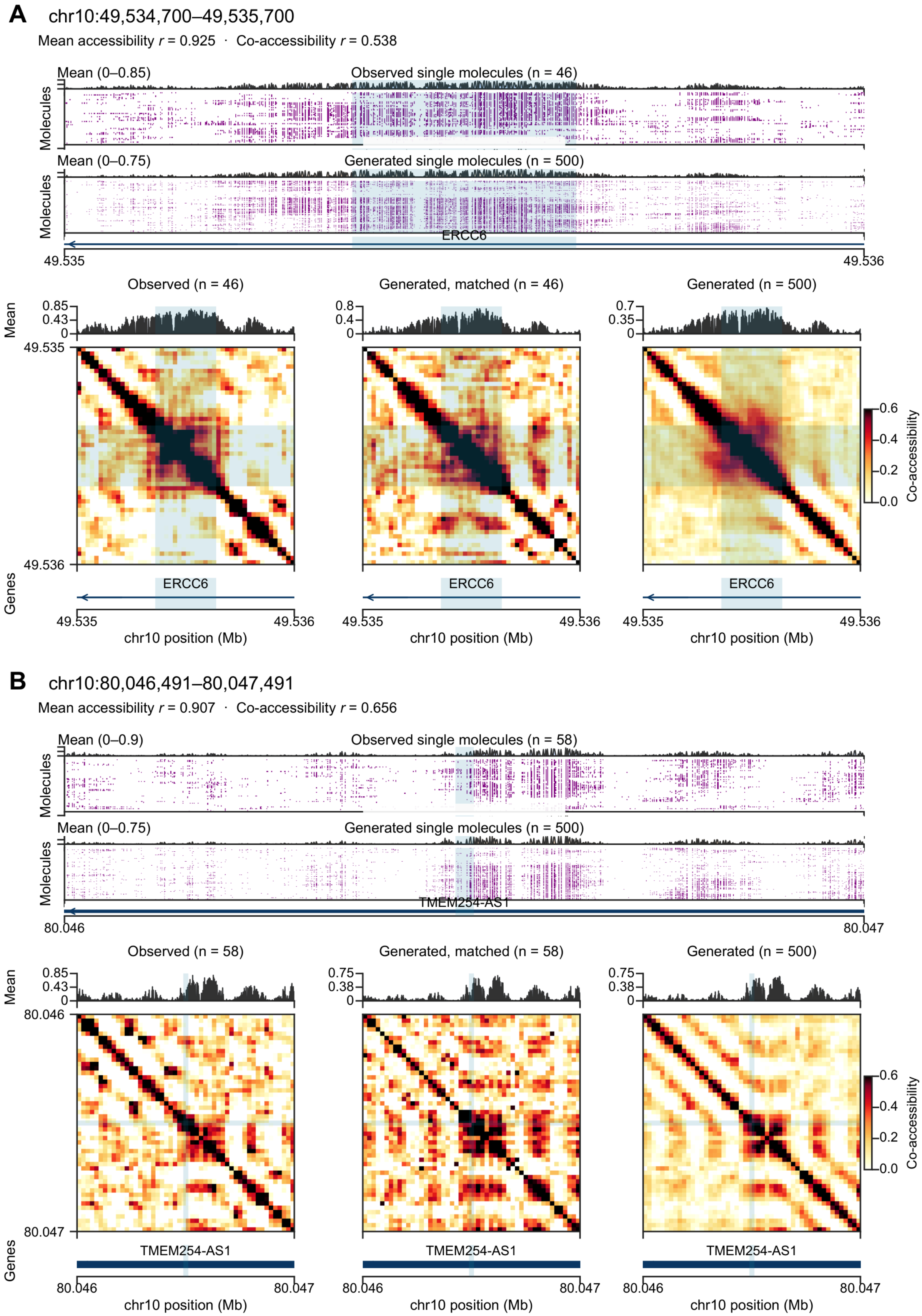


**Figure S5. Further held-out chromosome 10 examples.** As in Figure S3.

#### Figure S6

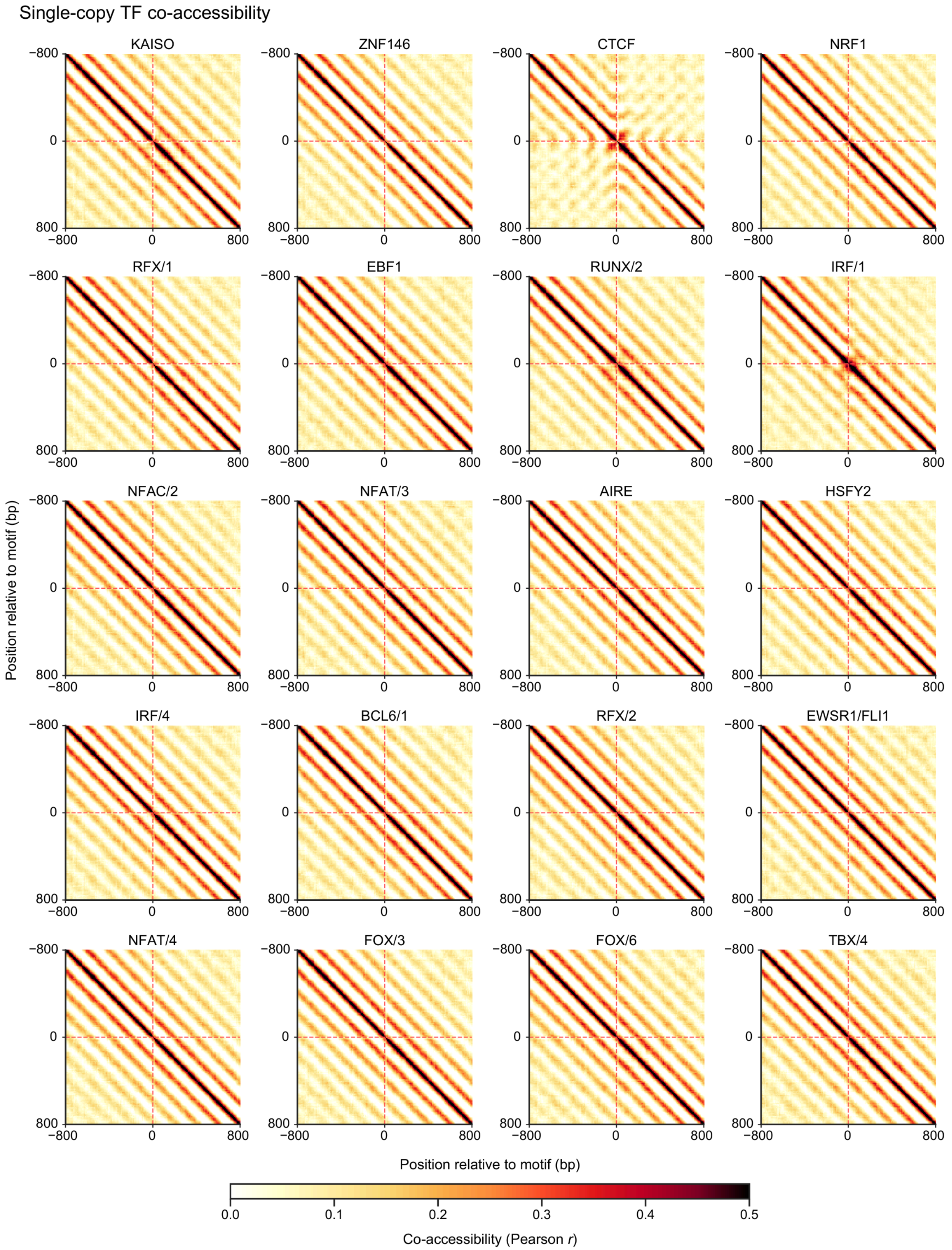


**Figure S6. Co-accessibility maps after single-copy motif insertion for 20 transcription factors.** Mean co-accessibility (20 bp bins, ±800 bp) of generated molecules after inserting one motif copy into 100 random backgrounds. Off-diagonal bands reflect the nucleosome array; CTCF, RUNX/2 and IRF/1 also show a block at the insertion.

#### Figure S7

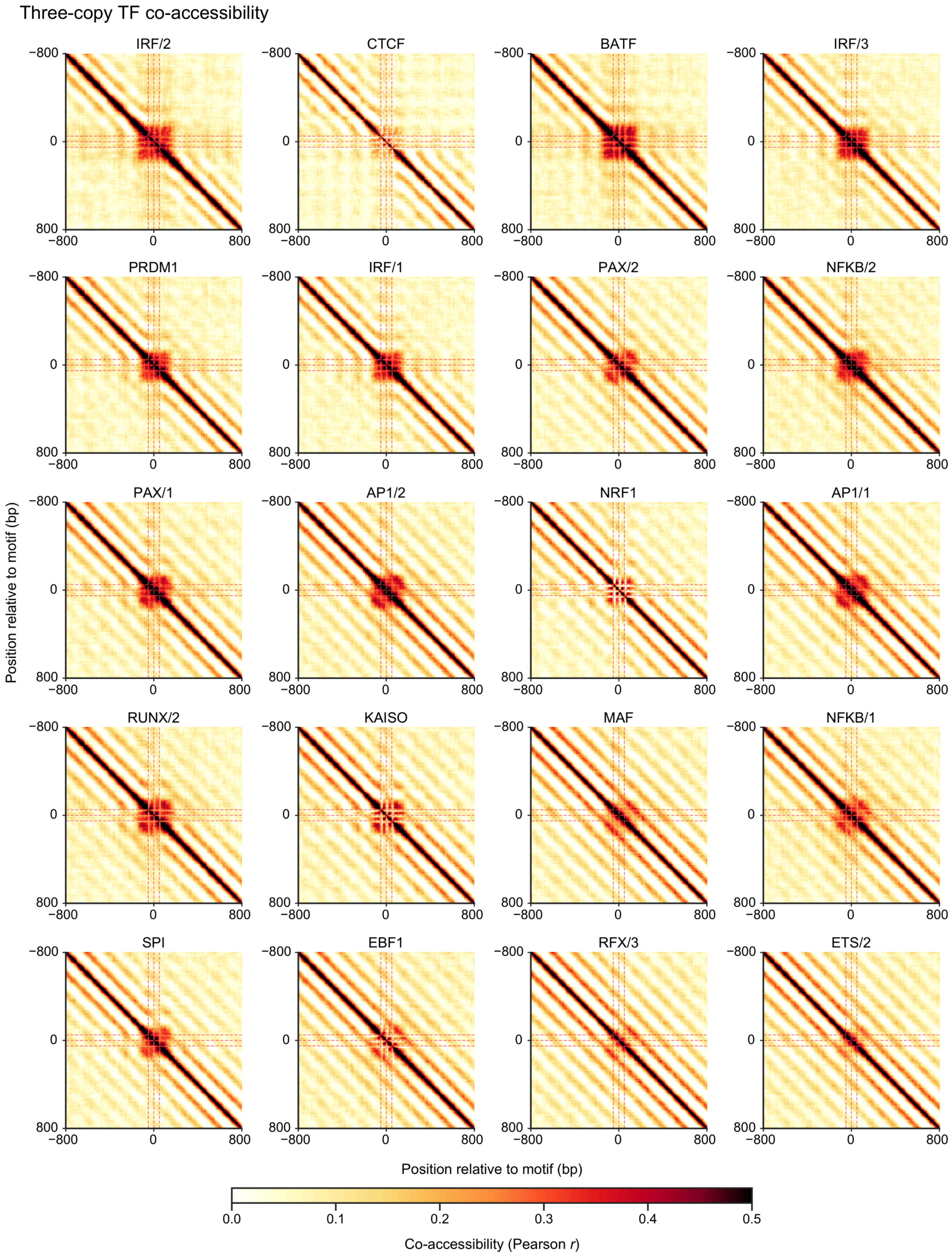


**Figure S7. Co-accessibility maps after three-copy motif insertion for 20 transcription factors.** As in Figure S6, with three copies 50 bp apart; motifs ordered by three-copy opening power.

#### Figure S8

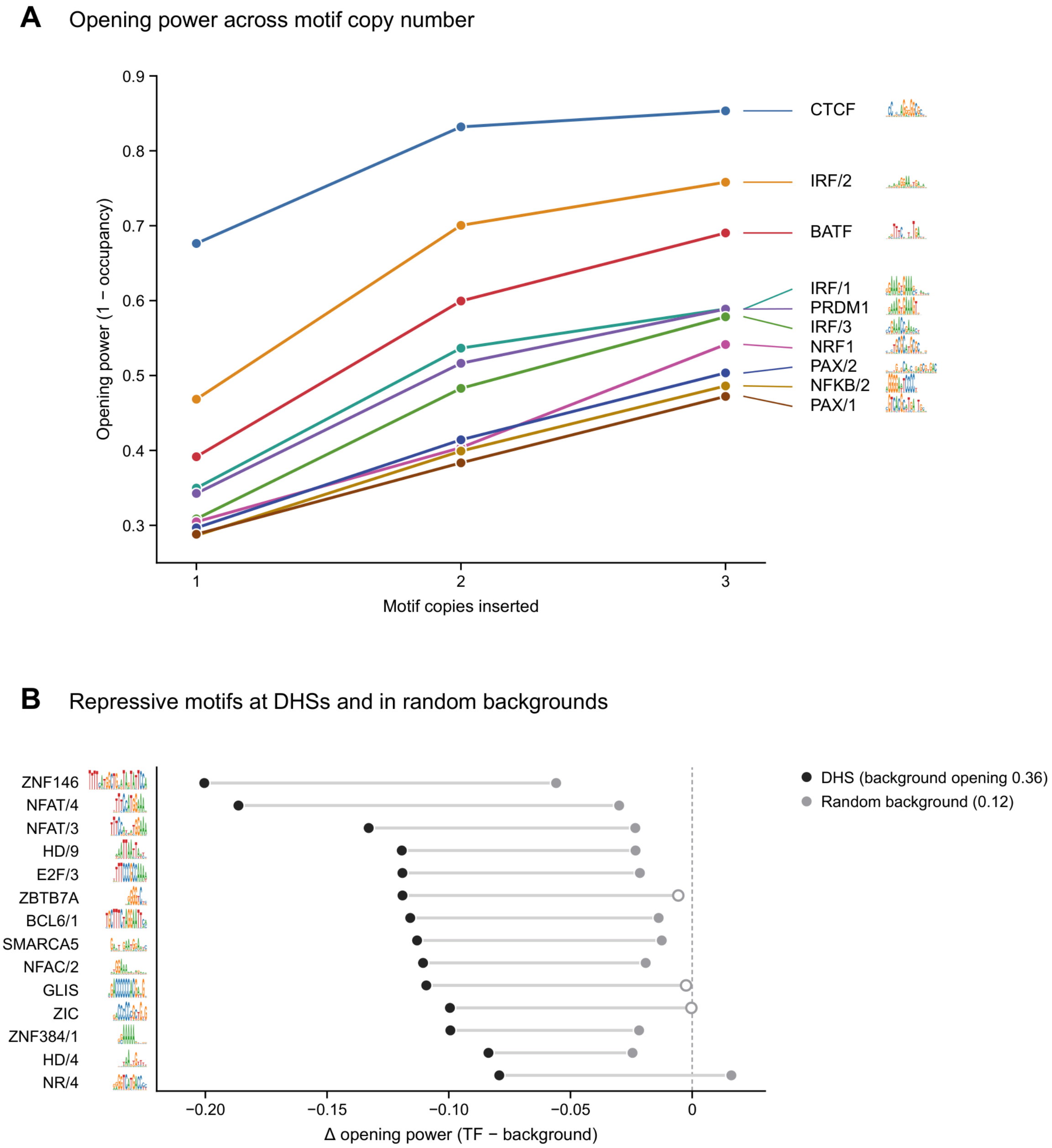


**Figure S8. Nucleosome opening increases with motif copy number.** (A) Opening power (1 − nucleosome occupancy at the inserted block) for one to three copies of the ten strongest three-copy motifs in 100 random backgrounds. (B) Change in opening power after one copy of repressive motifs at DHSs (black) or random backgrounds (grey); filled, Wilcoxon $q < 0.05$; open, not significant.

#### Figure S9

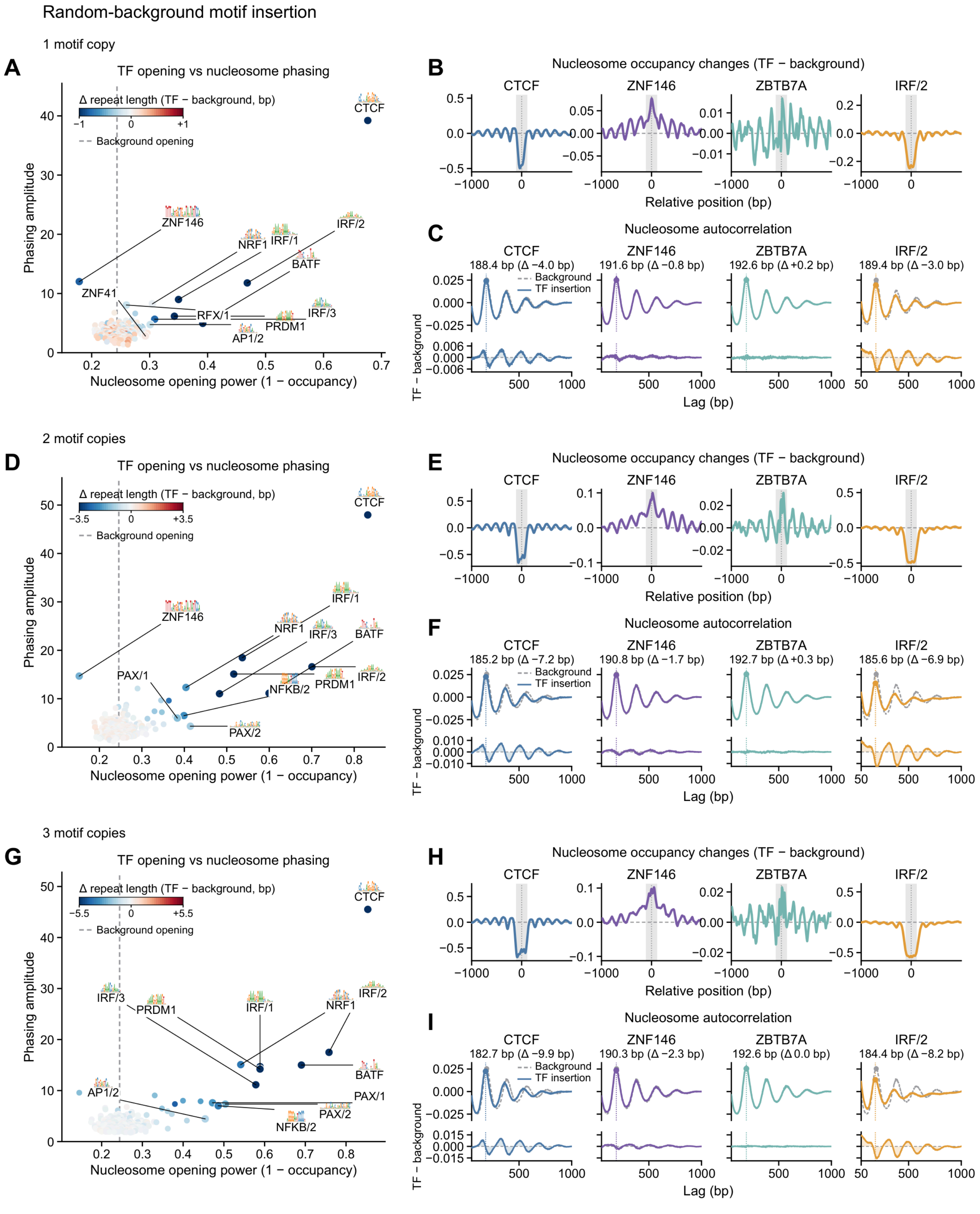

Random-background motif insertion
1 motif copy
A
TF opening vs nucleosome phasing
Δ repeat length (TF − background, bp)
Background opening
Phasing amplitude
Nucleosome opening power (1 − occupancy)
CTCF
ZNF146
NRF1
IRF/1
IRF/2
BATF
ZNF41
RFX/1
IRF/3
PRDM1
AP1/2
B
Nucleosome occupancy changes (TF − background)
CTCF
ZNF146
ZBTB7A
IRF/2
Relative position (bp)
C
Nucleosome autocorrelation
188.4 bp (Δ −4.0 bp)
191.6 bp (Δ −0.8 bp)
192.6 bp (Δ +0.2 bp)
189.4 bp (Δ −3.0 bp)
Background
TF insertion
TF − background
Lag (bp)
2 motif copies
D
TF opening vs nucleosome phasing
PAX/1
NFKB/2
PAX/2
E
Nucleosome occupancy changes (TF − background)
F
Nucleosome autocorrelation
185.2 bp (Δ −7.2 bp)
190.8 bp (Δ −1.7 bp)
192.7 bp (Δ +0.3 bp)
185.6 bp (Δ −6.9 bp)
3 motif copies
G
TF opening vs nucleosome phasing
H
Nucleosome occupancy changes (TF − background)
I
Nucleosome autocorrelation
182.7 bp (Δ −9.9 bp)
190.3 bp (Δ −2.3 bp)
192.6 bp (Δ 0.0 bp)
184.4 bp (Δ −8.2 bp)

**Figure S9. Opening, phasing, and nucleosome array structure across motif copy number.** Rows: one (A–C), two (D–F) and three (G–I) inserted copies in random backgrounds. (A, D, G) Opening power versus phasing amplitude for 285 motifs, colored by change in nucleosome repeat length; grey, below the noise floor (dotted). (B, E, H) Change in nucleosome occupancy for four motifs. (C, F, I) Mean single-molecule autocorrelation (dashed, background), with repeat length above and difference below.

#### Figure S10

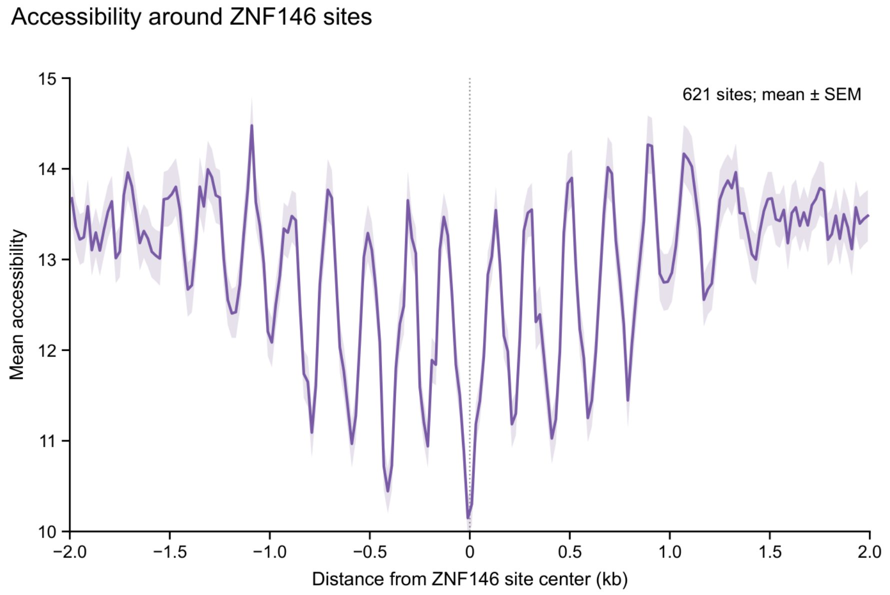


**Figure S10. Fiber-seq accessibility around endogenous ZNF146 sites.** Mean accessibility of GM12878 Fiber-seq molecules around 621 chromosome 10 ZNF146 ChIP-seq peaks; shading, s.e.m. The profile matches the nucleosome gain and phasing after ZNF146 insertion (Figure S9).

#### Figure S11

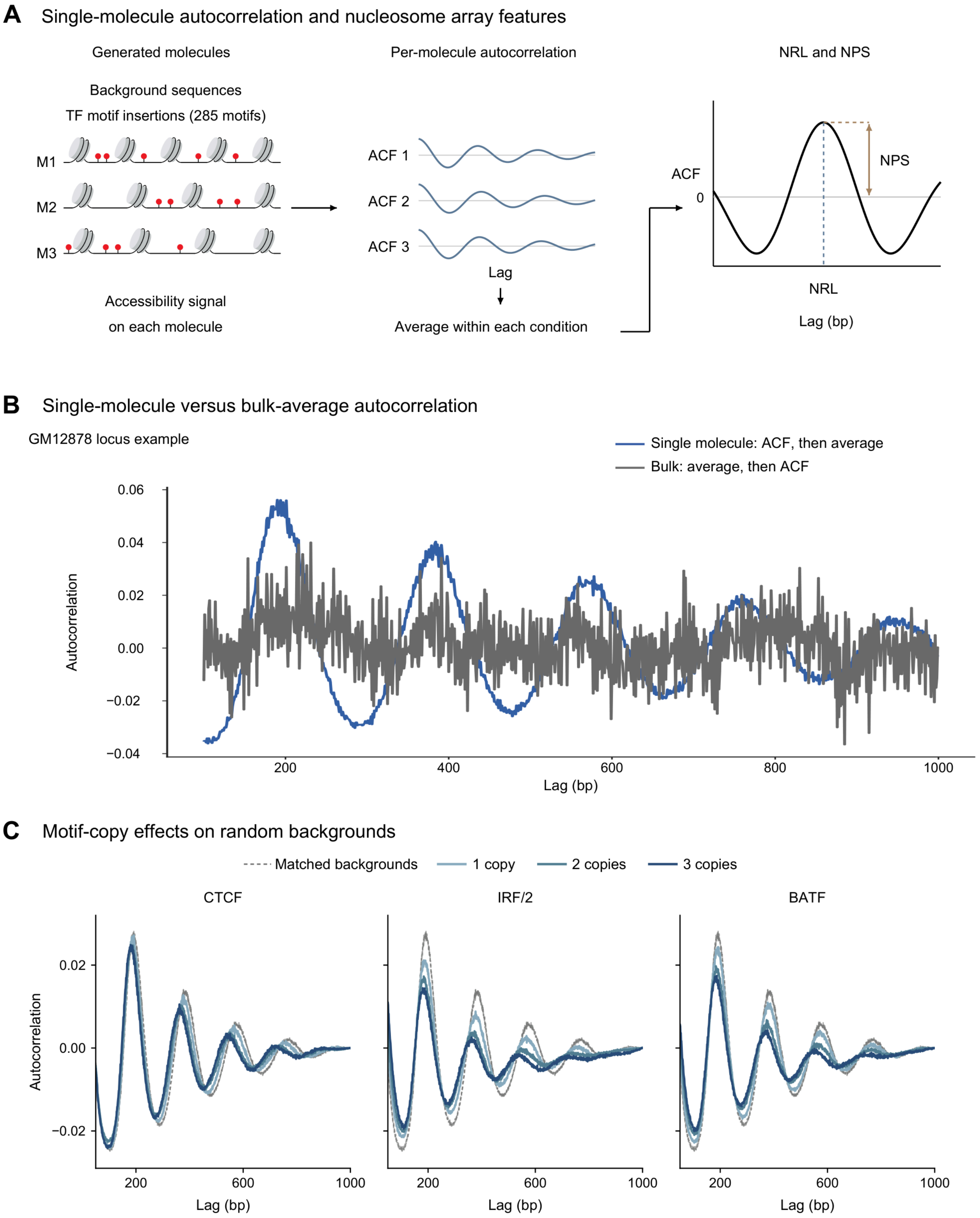


**Figure S11. Single-molecule autocorrelation as a readout of nucleosome array structure.** (A) Molecules are autocorrelated individually and the curves averaged; nucleosome repeat length (NRL) and periodicity strength (NPS) are the lag and height of the periodic peak. Schematic. (B) At a GM12878 CTCF locus, periodicity appears in the mean of single-molecule autocorrelations (blue) but not in the autocorrelation of the mean profile (grey). (C) Mean single-molecule autocorrelation after one to three inserted copies (light to dark blue); grey dashed, backgrounds.

#### Figure S12

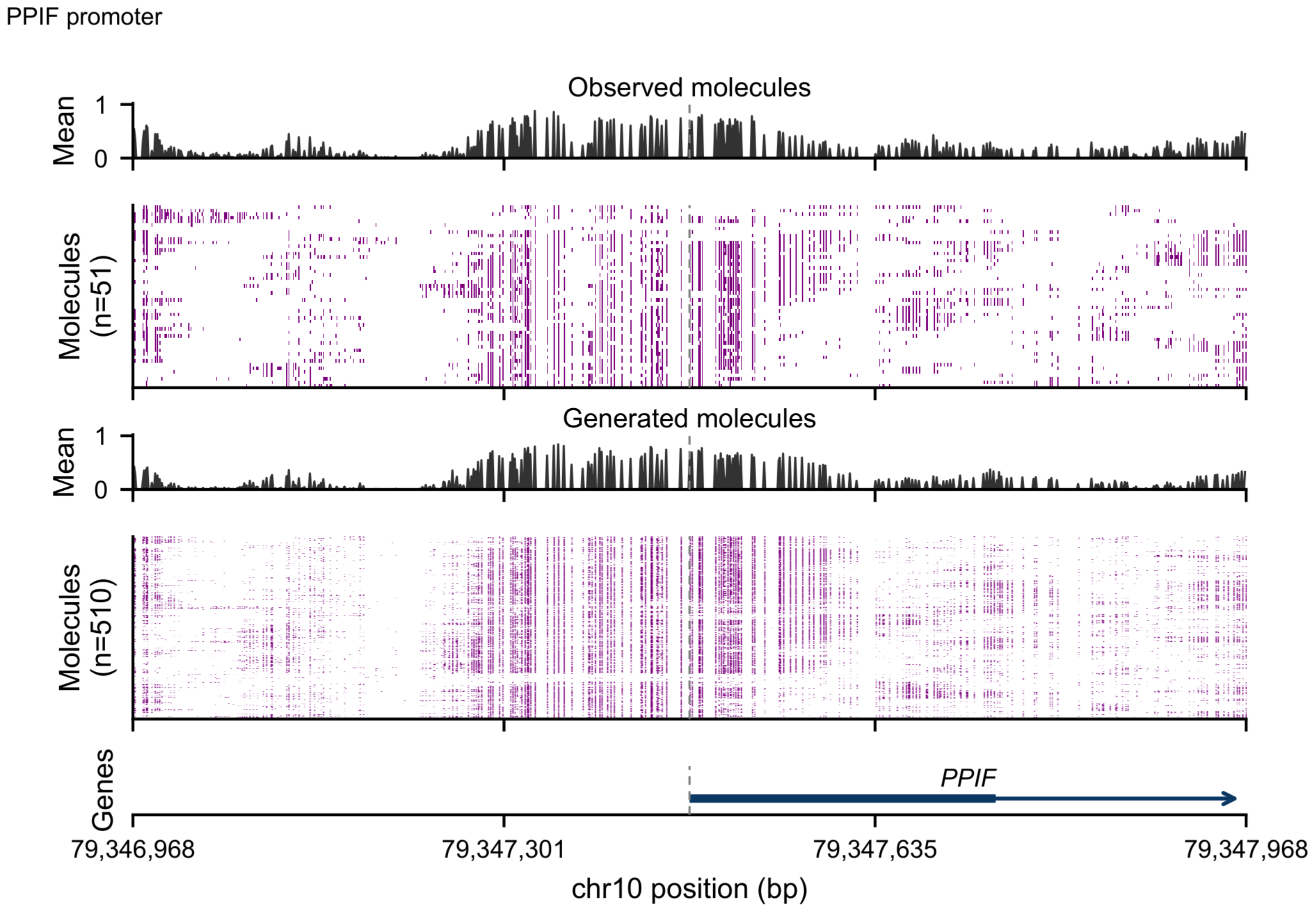


**Figure S12. Observed and generated chromatin at the PPIF promoter, a TATA-driven TSS on a held-out chromosome.** Single-molecule accessibility over 1 kb centered on the PPIF TSS; rows clustered. Both observed and generated molecules show a broadly accessible promoter without a well-positioned +1 nucleosome.

#### Figure S13

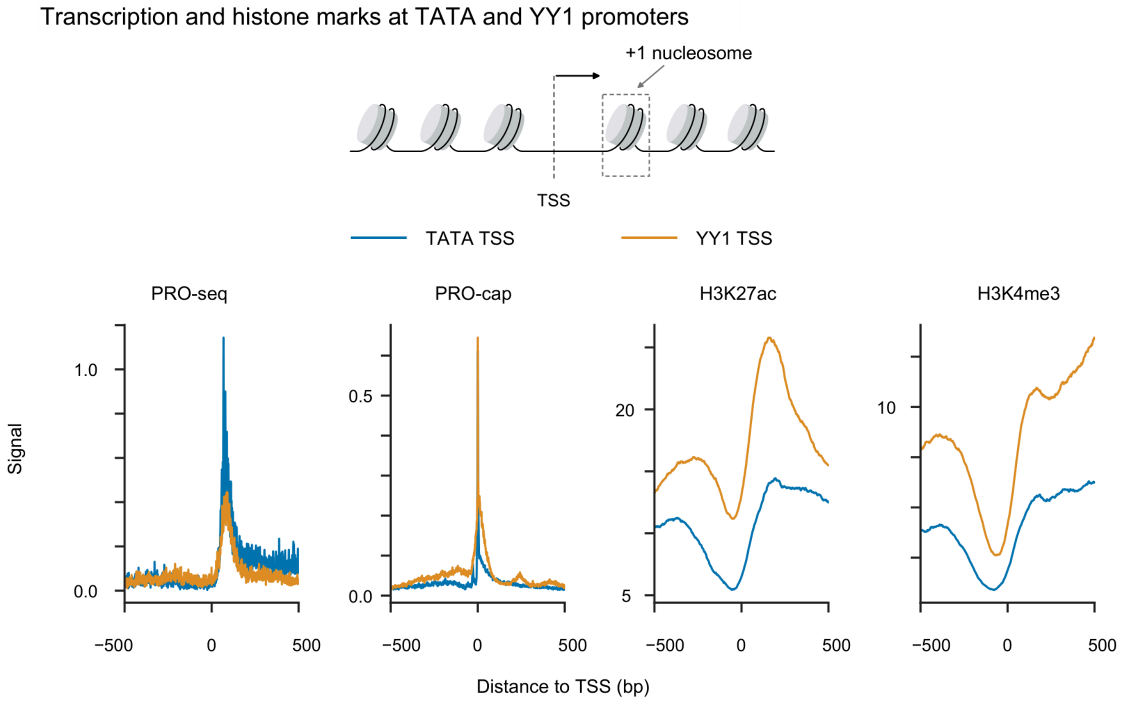


**Figure S13. Transcription and histone marks at TATA and YY1 promoters.** Mean PRO-seq, PRO-cap, H3K27ac and H3K4me3 signal at TATA-driven (blue) and YY1-driven (orange) TSSs. YY1 TSSs carry more H3K27ac and H3K4me3 and less PRO-seq signal at similar PRO-cap signal.

**Figure S14**

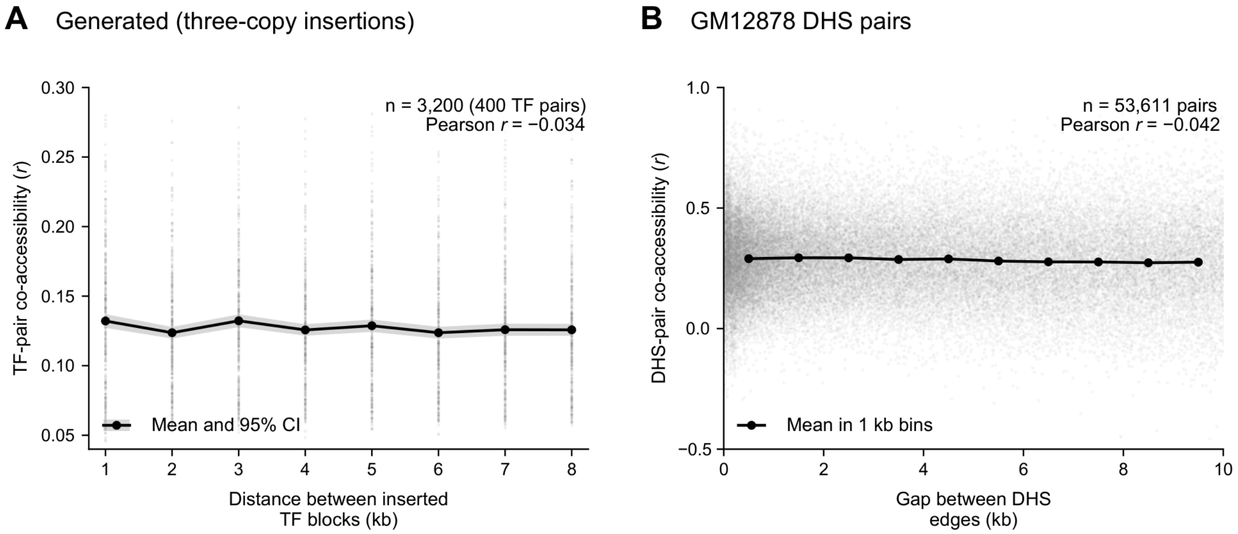


**Figure S14. Co-accessibility between distal elements depends little on their separation.** Co-accessibility versus separation for (A) pairs of inserted three-copy motif elements in generated molecules and (B) GM12878 DHS pairs in observed molecules. Grey, individual observations; black, means. Distances are defined differently, so only trends are comparable.

#### Figure S15

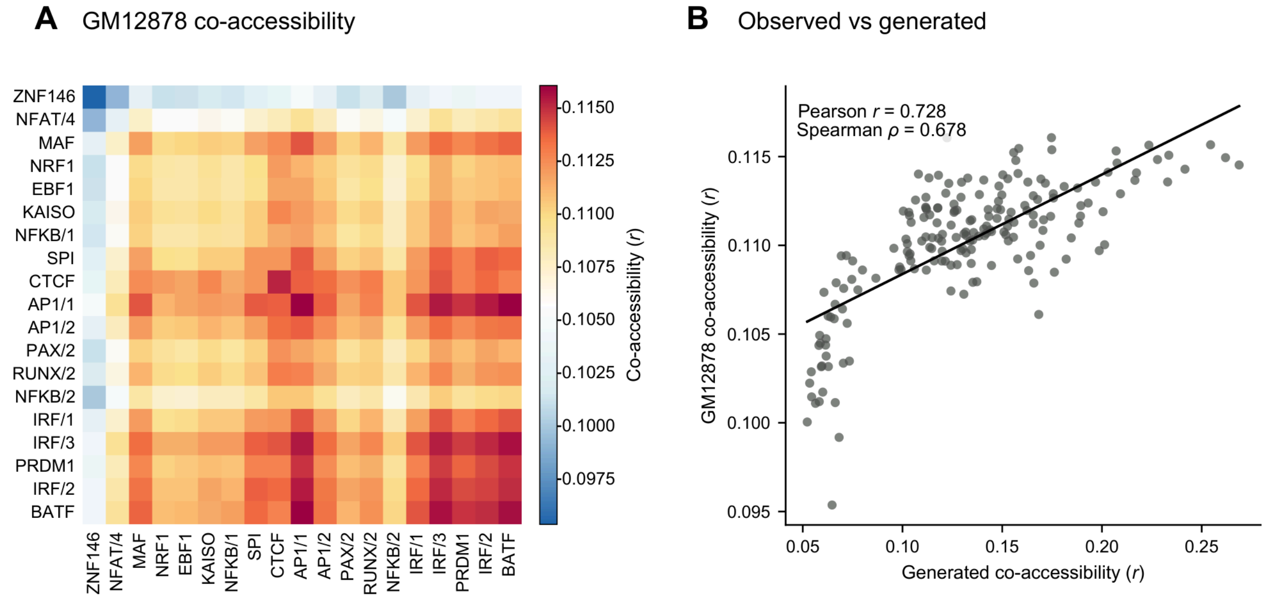


**Figure S15. Endogenous Fiber-seq co-accessibility agrees with the insertion model.** (A) TF-by-TF co-accessibility in GM12878 Fiber-seq molecules for 19 motifs (site pairs 1–10 kb apart, per-locus means removed), ordered as in Figure 4. (B) The same 190 TF pairs against three-copy insertion co-accessibility; line, least-squares fit.

#### Figure S16

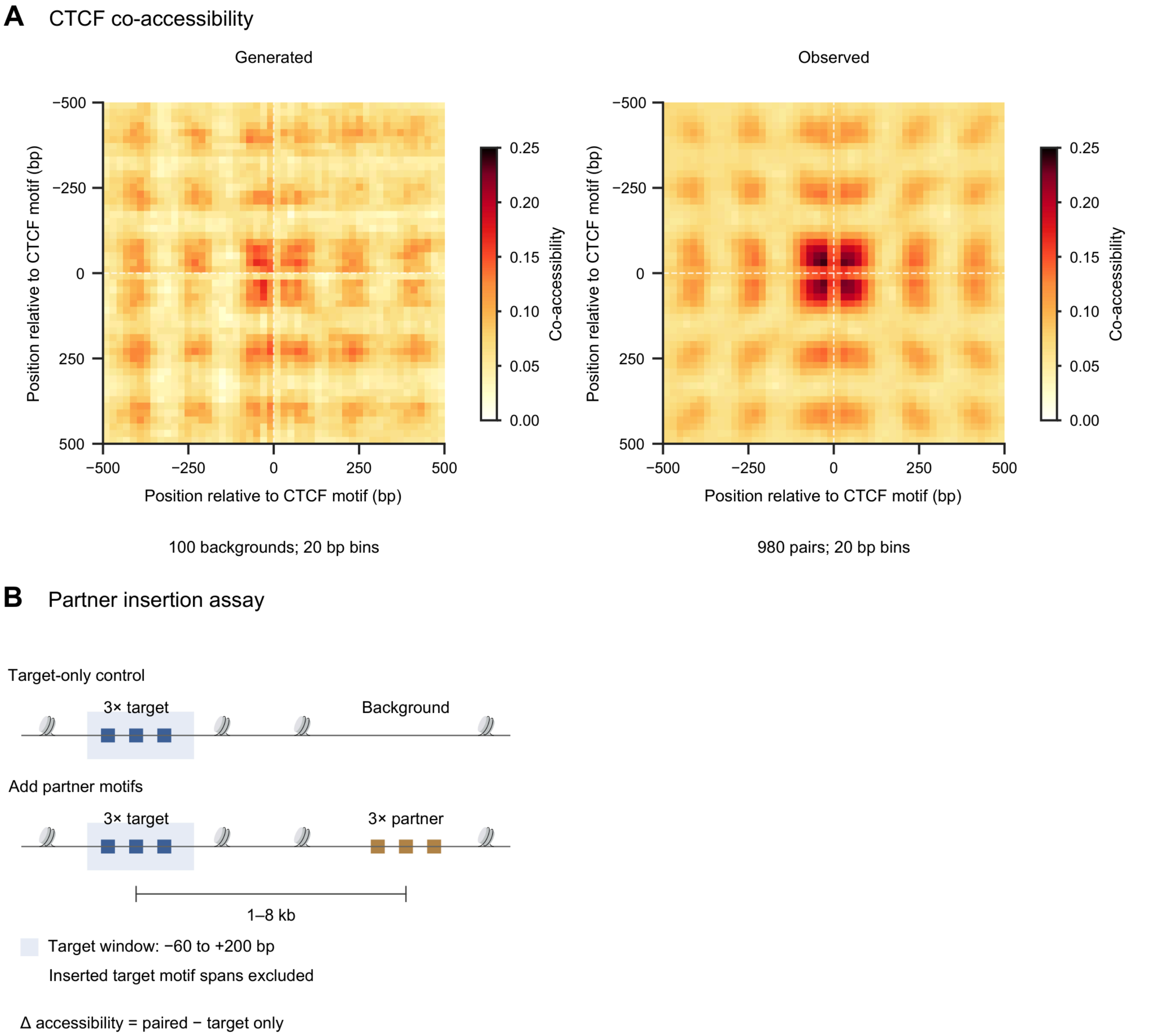


**Figure S16. CTCF co-accessibility and the partner-insertion assay.** (A) Co-accessibility around CTCF motifs in generated and observed molecules. (B) Partner-insertion assay: three target copies are inserted alone or with three partner copies 1–8 kb away; the change in target accessibility relative to the target-only control gives the matrix in Figure 4H. Schematic.

#### Figure S17

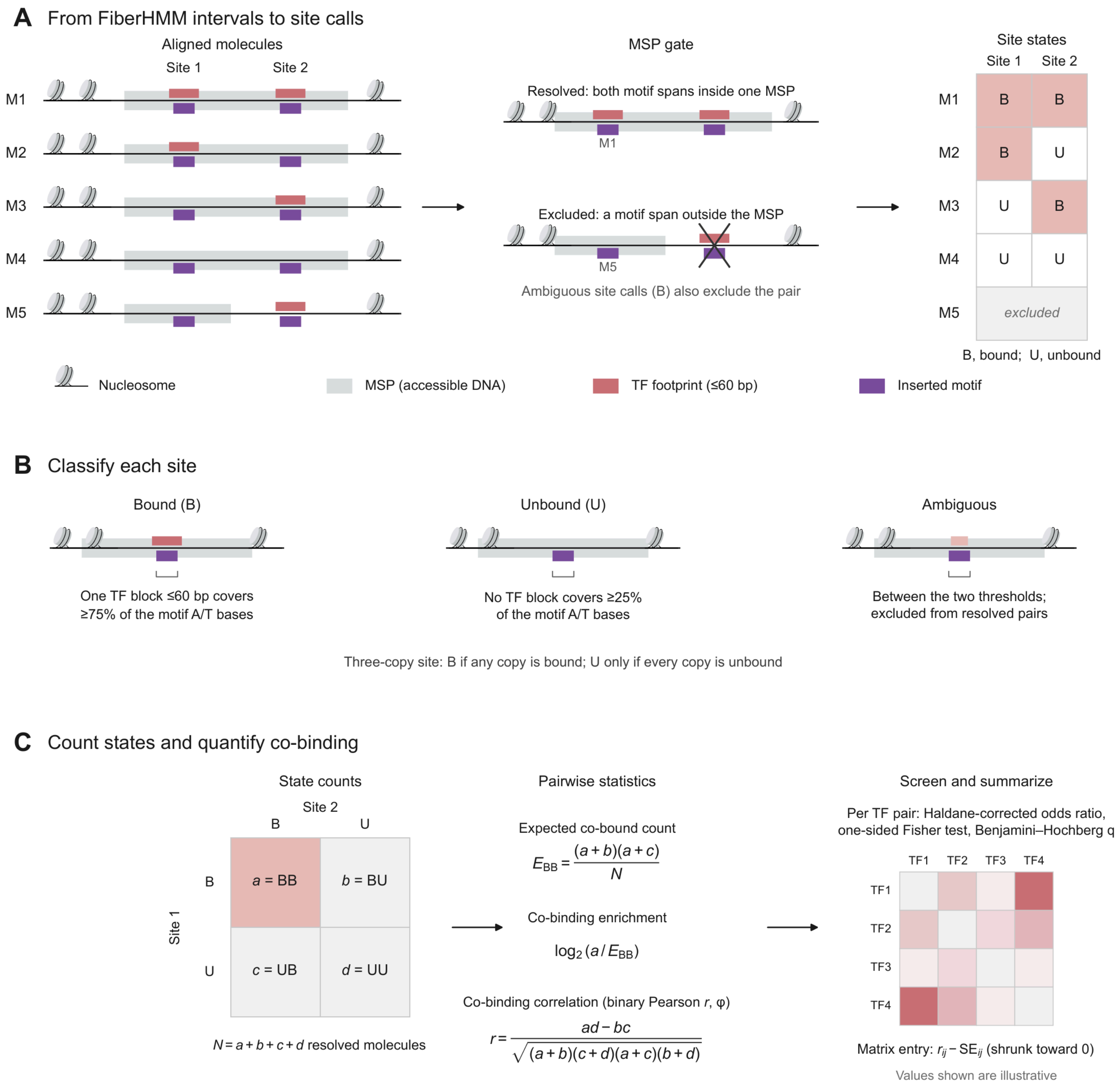


**Figure S17. Calling transcription factor co-binding on individual molecules with FiberHMM.** (A) FiberHMM calls nucleosomes, TF footprints (red, ≤60 bp) and methyltransferase-sensitive patches (MSPs; grey) on each generated molecule; a site pair is resolved only when both motif spans (purple) lie within one MSP. (B) A site is bound if one TF block covers ≥75% of its motif A/T bases and unbound if none covers ≥25%; other sites are excluded. (C) Resolved molecules form a 2 × 2 table of site states, giving co-binding enrichment and the binary co-binding correlation (φ). Values are illustrative.

#### Figure S18

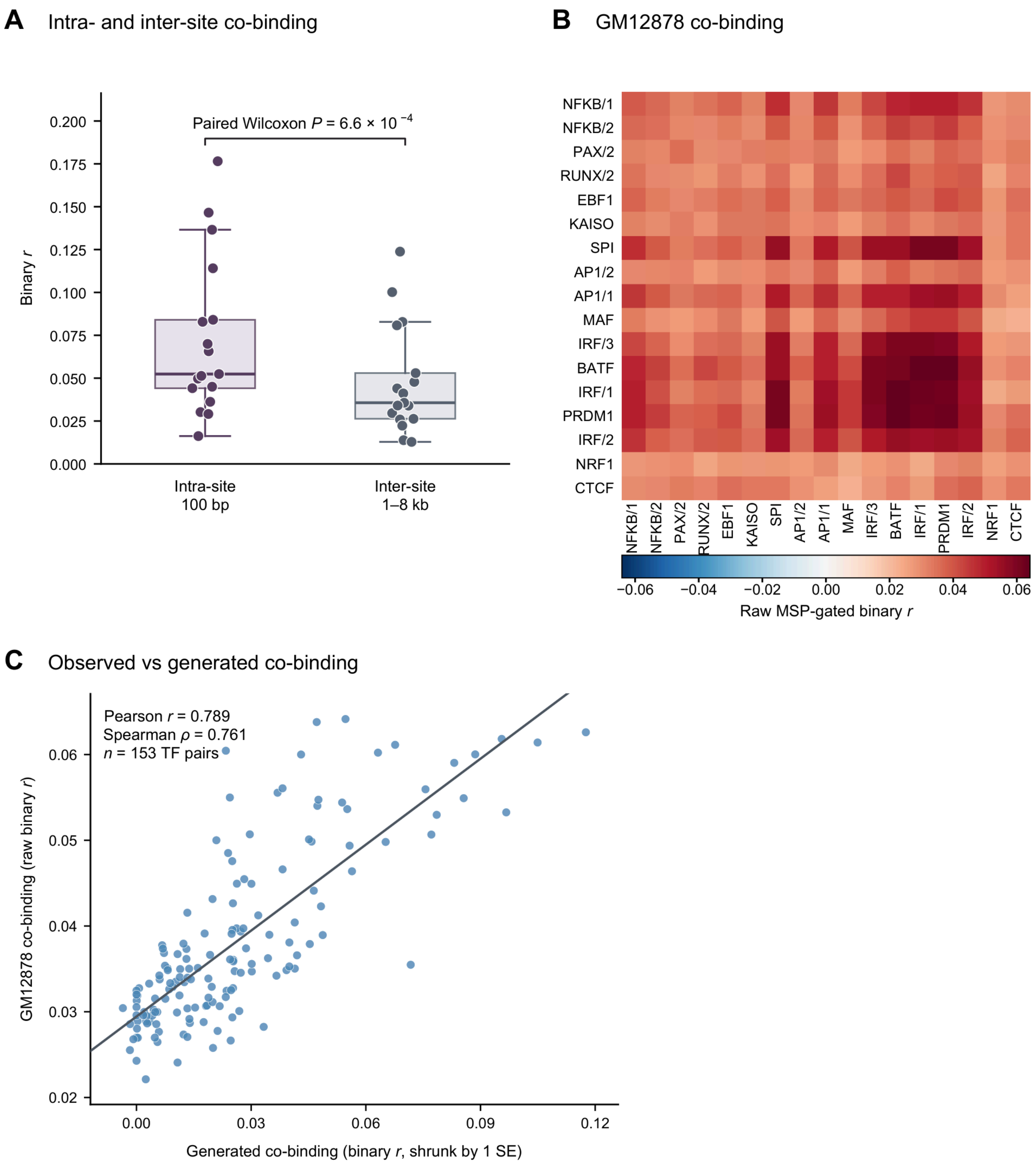


**Figure S18. Co-binding within and between elements, and agreement with Fiber-seq data.** (A) Binary co-binding correlation for 17 TF families with two homotypic copies 100 bp apart in one element (intra-site) or 1–8 kb apart (inter-site); boxes, median and interquartile range; whiskers, 1.5× interquartile range. (B) MSP-gated binary co-binding correlation in GM12878 Fiber-seq molecules. (C) Observed (B) versus generated co-binding, the latter shrunk by one standard error.

### Figure S19

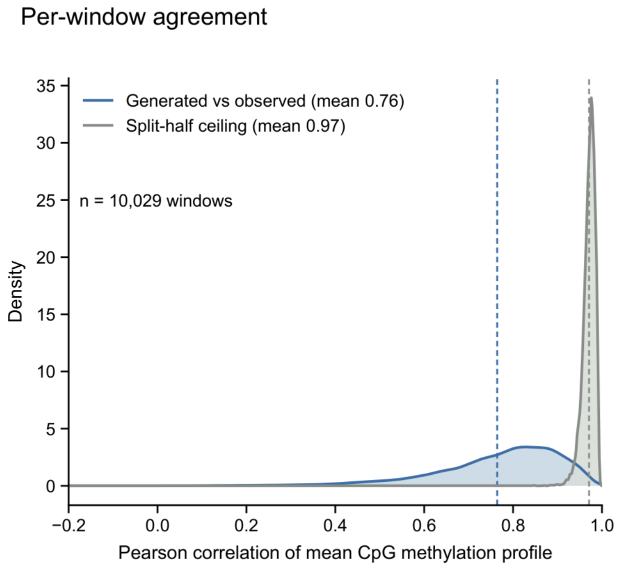


**Figure S19. Accuracy of predicted mean CpG methylation on held-out chromosomes.** Correlation between generated and observed mean CpG methylation profiles (blue) and its split-half ceiling (grey) across held-out 10 kb DHS windows (chromosomes 8–10); dashed lines, means. Pooled efficiency (mean correlation divided by mean ceiling) is 79% (95% bootstrap interval 78–79%).

#### Figure S20

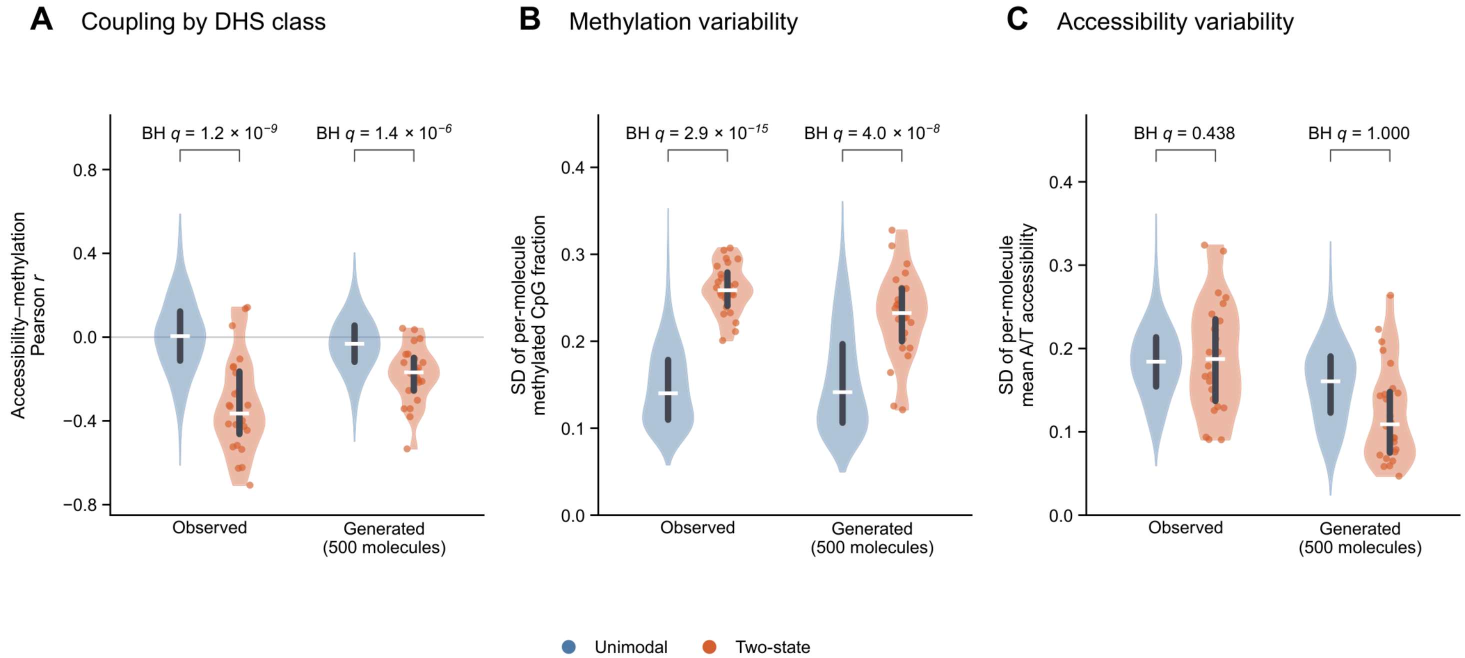


**Figure S20. Accessibility–methylation coupling and molecule-level variability by DHS class.** Held-out native DHSs on chromosomes 8–10 classed as unimodal (grey; n = 2,718) or two-state (orange; n = 28) by beta-binomial mixture modeling, for observed and 500 generated molecules per DHS; white bars, medians; dark bars, interquartile ranges; one-sided Mann–Whitney tests, Benjamini–Hochberg-adjusted. (A) Cross-molecule accessibility–methylation correlation. (B, C) Standard deviation across molecules of methylated-CpG fraction (B) and mean A/T accessibility (C).

#### Figure S21

Motifs enriched in two-state DHSs

| Motif | | *P* |
|---|---|---|
| NRF1 | 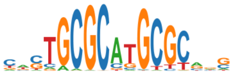 | $6.9 \times 10^{-5}$ |
| ZBTB14 | 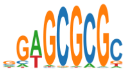 | $1.3 \times 10^{-4}$ |

**Figure S21. Motifs enriched in two-state DHSs.** Two-state DHSs were matched within chromosome to unimodal DHSs on CpG count, mean methylation, mean accessibility and molecule depth, and scores for 268 motif archetypes were compared by one-sided paired Wilcoxon tests. Shown are the two archetypes significant after Benjamini–Hochberg correction (q < 0.05), with their P values.

#### Figure S22

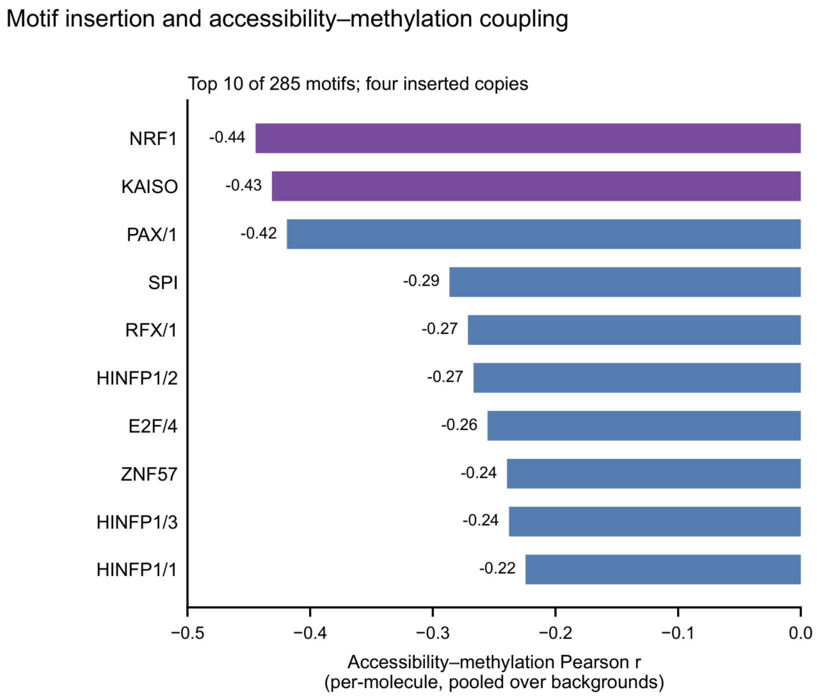


**Figure S22. Motif insertion and accessibility–methylation coupling.** Correlation between per-molecule accessibility and CpG methylation after inserting four copies of each motif into 20 backgrounds (100 generated molecules each), for the ten strongest of 285 motifs; purple, CpG-rich motifs.

#### Figure S23

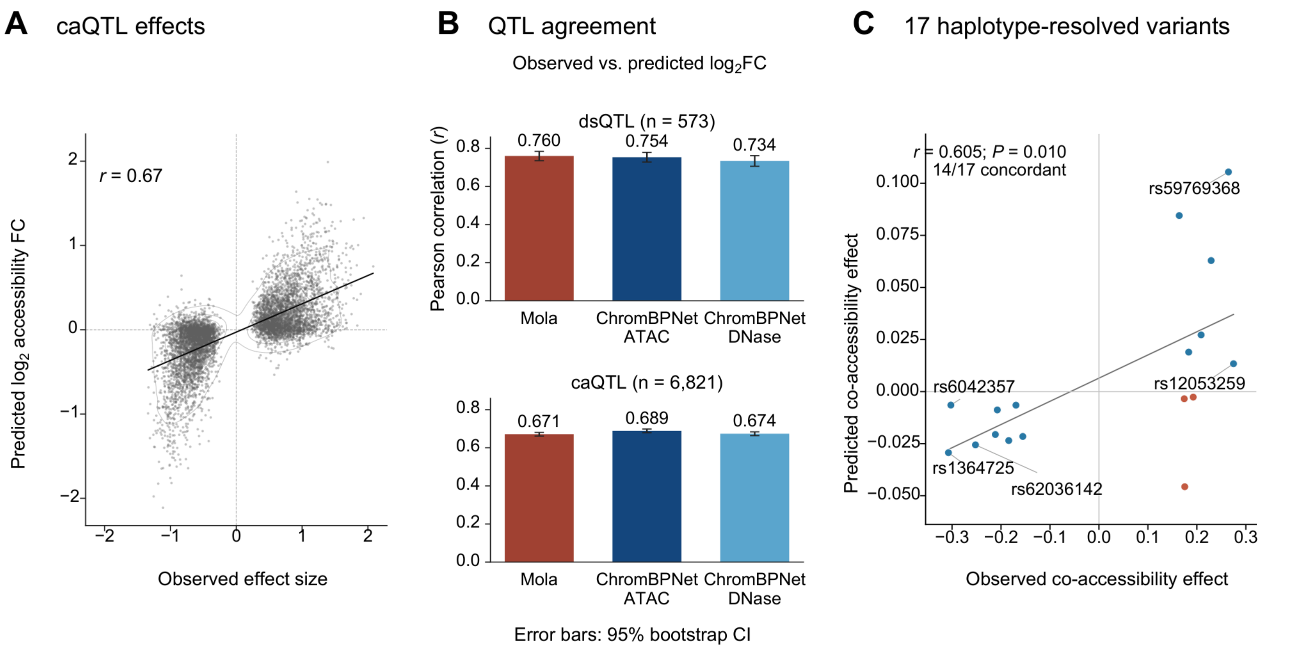


**Figure S23. Variant effects on mean accessibility and on distal co-accessibility.** (A) Predicted versus measured log2 accessibility fold change for African lymphoblastoid caQTLs; line, least-squares fit. (B) Correlation with measured effects for Mola and ChromBPNet (ATAC- and DNase-trained) in dsQTLs and caQTLs; error bars, 95% bootstrap CI. (C) Measured versus predicted allele-specific co-accessibility between the variant window and the neighboring DHS for 17 haplotype-resolved GM12878 variants; blue, concordant; red, discordant.
